# Effect of leaflet asymmetry on the stretching elasticity of lipid bilayers with phosphatidic acid

**DOI:** 10.1101/2022.10.12.511874

**Authors:** Dominik Drabik, Piotr Hinc, Mareike Stephan, Rafaela R. M. Cavalcanti, Aleksander Czogalla, Rumiana Dimova

## Abstract

The asymmetry of membranes has a significant impact on their biophysical characteristics and behavior. This study investigates the composition and mechanical properties of symmetric and asymmetric membranes in giant unilamellar vesicles (GUVs) made of phosphatidylcholine (POPC) and phosphatidic acid (POPA). A combination of fluorescence quantification, zeta potential measurements, micropipette aspiration and bilayer molecular dynamics simulations are used to characterize these membranes. The outer leaflet composition in vesicles is found consistent across the two preparation methods we employed, namely electroformation and inverted emulsion transfer. However, characterizing the inner leaflet poses challenges. Micropipette aspiration of GUVs show that oil residues do not substantially alter membrane elasticity, but simulations reveal increased membrane thickness and decreased interleaflet coupling in the presence of oil. Asymmetric membranes with a POPC:POPA mixture in the outer leaflet and POPC in the inner leaflet display similar stretching elasticity values to symmetric POPC:POPA membranes, suggesting potential POPA insertion into the inner leaflet during vesicle formation and suppressed asymmetry. The inverse compositional asymmetry, with POPC in the outer leaflet and POPC:POPA in the inner yield less stretchable membranes with higher compressibility modulus compared to their symmetric counterparts. Challenges in achieving and predicting compositional correspondence highlight the limitations of phase-transfer-based methods. Additionally, caution is advised when using fluorescently labeled lipids (even at low fractions of 0.5 mol%), as unexpected gel-like domains in symmetric POPC:POPA membranes were observed only with a specific type of labeled DOPE (dioleoylphosphatidylethanolamine) and the same fraction of unlabeled DOPE. The latter suggest that such phase separation may result from interactions between lipids and membrane fluorescent probes. Overall, this study underscores the complexity of factors influencing GUV membrane asymmetry, emphasizing the need for further research and improvement of characterization techniques.

**SIGNIFICANCE:** Asymmetrically charged lipid bilayer models are superior to commonly used symmetrical ones, exhibiting naturally present asymmetry, thereby exhibiting a more adequate range of biophysical membrane characteristics better reflecting biological membranes. This study focuses on the mechanical properties of phosphatidic acid (PA)-enriched membranes, a crucial lipid for cellular lipid metabolism, e.g. glycerophospholipid synthesis, and for signal transduction. **M**icropipette aspiration, fluorescent PA-sensor, and zeta potential studies demonstrate that asymmetric membranes are less stretchable than symmetric ones. Accompanying *in silico* studies on the symmetric membranes confirm that oil impurities do not influence the membrane stretching elasticity but increase its thickness and decrease the coupling of the two leaflets, which sheds light on the elastic behavior of experimental models of asymmetric lipid bilayers.

## 1. INTRODUCTION

Lipid vesicle models are commonly used to study the behavior and dynamics in biological membranes. Among them, giant vesicles represent a convenient system to directly explore and visualize the membrane under a microscope [1-4]. As simplified biomembranes, often formed with a spherical shape, they can be employed to assess various membrane characteristics related to lipid dynamics, mechanical or electrical properties and organization. For instance, giant vesicles have been successfully used for cell function studies [5-7], as artificial tissue models [8] and for resolving protein-lipid interactions [9, 10], to list just a few. However, one significant difference between biological membranes and those of synthetic (or even cell-derived) giant vesicles as investigated in a large bulk of studies is the lack of asymmetry in the latter. In living cells, membrane asymmetry is a prevalent and fundamental feature [11-13]. While lipidomics studies have shown the presence of a variety of lipids within different cellular membranes, the significance of their transversal distribution remains poorly understood. The biophysical consequences of membrane asymmetry and the associated functional significance represent an emerging topic in cell physiology [14]. Asymmetry plays a crucial role in the function of membrane proteins and lipid molecules, particularly those exclusively present in or associated with one of the membrane leaflets. By studying the distribution and dynamics of these lipids, we can gain valuable insights into the intricate mechanisms that govern cellular membranes and their interactions with the environment.

One such lipid type is phosphatidic acids (PAs). PAs have a negative charge and a phosphomonoester group. Compared to other membrane lipids, they are characterized by a relatively small head group. They are essential components in lipid metabolism, serving as precursors to the synthesis of various glycerophospholipids and modulating the shape and curvature of cellular membranes [15, 16]. PAs have been also proven to act as potent signaling molecules that govern several important aspects of cell biology [17]:, e.g. regulate activity of key metabolism-orchestrating kinase mTOR [10, 18]. Symmetric models of phosphatidylcholine (PC) membranes containing various types of PA have not shown particularly different characteristics (such as bending rigidity and area per lipid) that are specific to the PA lipid type [19]. However, in biological systems, PAs have been observed to localize primarily within the inner leaflet of plasma membranes [11, 20], and the enzymes responsible for PA synthesis, such as phospholipases D diacylglycerol kinase, are found to operate only at one of the leaflets of the plasma membrane and subcellular compartments [15, 21, 22]. Thus, the characterization of asymmetric model systems, and comparison to their symmetric counterparts, could provide important information related to PA synthesis and membrane incorporation. The preparation of asymmetric models with PA in vesicles with sizes in the 100 nm range has already been reported [23]. In this work, we use cell-sized giant unilamellar vesicles (GUVs) and our focus is on assessing the mechanical properties of such membranes, which could provide valuable insights into how asymmetry affects the membrane stability, elasticity, and other physical properties.

Progress towards understanding asymmetric membranes using giant vesicles became possible after the development of methods that allow the preparation of synthetic asymmetric membranes models [1, 4]. The most common and still most-widely used method for production of GUVs with asymmetric leaflets is the phase-transfer method (also known as oil-droplet or emulsion transfer, or inverted emulsion method), first introduced by Träuble and Grell [24] and later employed in various studies [25-29]. The approach involves the formation of each leaflet separately, followed by their combination under external force such as gravity, centrifugation or microfluidic flow. The first step consists of the preparation of water-in-oil emulsion where the aqueous phase is dispersed in a nonpolar solvent (e.g. oil) containing lipids, which form a monolayer at the water-oil interface of the emulsion droplets. The droplets are then covered by a second lipid monolayer. This is achieved by pulling them through another oil-water interface, stabilized by the lipid building the outer leaflet, to obtain the vesicle architecture. Alternative approaches to generate GUVs with asymmetric membrane composition are based on leaflet lipid exchange via hemifusion [30], cyclodextrin-mediated lipid exchange [31, 32] and microfluidic jetting [33, 34]. These approaches have been applied predominantly on phosphatidylcholine-based membranes and it is questionable whether the use of other types of lipids, for example with different amphiphilicity (or charge) and preference to the oil-water interface, would also lead to asymmetric membranes with compositional correspondence. Indeed, a previous study has demonstrated that cholesterol incorporates at a very low fraction in vesicles prepared using a phase-transfer approach [35].

There are only a few reports studying the mechanical properties of GUVs with asymmetric lipid membranes. These studies address the mechanics of POPC:DOPC (palmitoyloleoyl phosphatidylcholine:dioleoyl phosphatidylcholine) asymmetric vesicles obtained via the phase transfer method [36], reporting a bending rigidity almost twice higher than that of the corresponding symmetric systems. Similar effect was confirmed for asymmetric vesicles obtained with a microfluidic device [34]. Additionally, DMPC:DOPC (dimyristoyl phosphatidylcholine:dioleoyl phosphatidylcholine) vesicles obtained using custom-made microfluidic devices [37], were shown to exhibit increased bending rigidity and area compressibility of the investigated asymmetric system compared to the symmetric ones. Furthermore, membranes with higher charge asymmetry were shown to be more prone to destabilization when exposed to pore-inducing electric pulses, suggesting that asymmetrically charged membranes are less stable [38].

In this work, we investigate the mechanical properties of asymmetric POPC:POPA membranes, where POPA is targeted to either the inner or the outer leaflet. To eliminate any possible influence of the preparation approach, we compare the asymmetric GUV systems to symmetric ones obtained using two different methods. To avoid misleading conclusions due to uncertainty regarding the membrane composition, we employed various approaches for accurate membrane characterization such as the use of α-synuclein-mEGFP as a PA sensor. This protein has been shown to strongly bind to membranes containing PA [39, 40]. Using micropipette aspiration, we probe the stretching elasticity of the membranes and discuss the resolved differences between the symmetric versus the asymmetric system and the influence of oil residues in the membrane resulting from the use of the phase-transfer preparation method. Overall, our study provides valuable insights into the properties of asymmetric lipid membranes containing PA and sheds light on their role in biological systems.

## 2 MATERIALS AND METHODS

### 2.1 Materials

The lipids POPC (1-palmitoyl-2-oleoyl-sn-glycero-3-phosphatidylcholine), POPA (1-palmitoyl-2-oleoyl-sn-glycero-3-phosphatidic acid), DOPE (1,2-dioleoyl-sn-glycero-3-phosphoethanolamine) and fluorescent probes NBD-PC (1-palmitoyl-2-{6-[(7-nitro-2-1,3-benzoxadiazol-4-yl)amino]hexanoyl}-sn-glycero-3-phosphocholine), Rh-DOPE (1,2-dioleoyl-sn-glycero-3-phosphoethanolamine-N-(lissamine rhodamine B sulfonyl), ammonium salt) were purchased from Avanti Polar Lipids (Alabaster, AL, USA). The fluorescent probe TexasRed-DHPE (1,2-dihexadecanoyl-sn-glycero-3-phosphoethanolamine, triethylammonium salt) was purchased from Invitrogen (Waltham, MS, USA). Rh-DHPE (N-(Lissamine rhodamine B sulfonyl)-1,2-dihexadecanoyl-sn-glycero-3-phosphoethanolamine, triethylammonium salt) was purchased from Biotium (SF, USA). Fluorecent probe Atto488-DOPE (1,2-Dioleoyl-sn-glycero-3-phosphoethanolamine labeled with Atto-488) was purchased from Sigma-Aldrich (international). Sucrose, glucose, imidazole, TEMED (N,N,N′,N′-tetramethyl ethylenediamine), lysozyme from chicken egg white, Sigma-Aldrich light mineral oil (density 0.83 g/cm^3^,1L bottle), BCA Protein Assay Kit and oligonucleotides were purchased from Sigma Aldrich (international). Low melting temperature agarose was purchased from Fisher Bioreagents (Massachusetts, USA). Roth mineral oil (density 0.88 g/cm^3^, 10 mL bottle - large volumes were avoided to minimize issues with humidity during handling the oil solutions), IPTG (isopropyl-β-D-thiogalactopyranosid), TRIS (tris-(hydroxymethyl)-amino methane), PMSF (phenylmethyl sulphonyl fluoride), Triton® X-100, HEPES (N-2-hydroxyethylpiperazine-N’-2-ethane sulphonic acid) and Rotiphorese® Gel 30 were purchased from Roth (international). Restriction enzymes EcoRI-HF, BamHI-HF, DpnI, as well as Phusion® High-Fidelity DNA Polymerase Quick Ligation™ Kit, Q5® Site-Directed Mutagenesis Kit and NiCo21(DE3) Competent *E. coli* were purchased from New England Biolabs (Ipswich, MS, USA). Ultrapure dNTPs Mix, agarose electrophoresis grade and OMNI nuclease were purchased from EURx (Poland). NucleoSpin® plasmid isolation kit and NucleoSpin® gel and PCR clean-up kit were purchased from Macherey-Nagel (Germany). PierceTM protease inhibitor tablets EDTA free and Pierce™ high capacity Ni-IMAC resin were purchased from Thermo Scientific (international). TALON® Metal Affinity Resin was purchased from Takara Bio (Mountain View, CA, USA). Econo-Pac 10DG Desalting Columns and 4-20% gradient Mini-PROTEAN®TGX™ Precast Gels were purchased from Bio-Rad Laboratories (USA). Protein Labeling Kit RED-NHS 2^nd^ Generation was purchased from NanoTemper Technologies (München, Germany). Glycerol, sodium chloride and ammonium persulfate were purchased from POCH (Poland). LB Miller broth was purchased from IBI Scientific (international). Kanamycin sulfate was purchased from BioShop (Poland). Coomasie Brillat Blue R-250 was purchased from PanReac AppliChem (Germany). Spectra/Por 6 Dialysis Tubing 25kD was purchased from Spectrum Laboratories (USA). pET3a aSyn murine plasmid containing a gene encoding mouse α-synuclein was a gift from Gabriele Kaminski Schierle (Addgene plasmid # 108865; http://n2t.net/addgene:108865; RRID:Addgene_108865).

### 2.2 Vesicle preparation

Both asymmetric and symmetric vesicles were obtained using the inverted emulsion method (also known as the phase-transfer method) [41-43] with some modifications. This method requires dissolving the lipids in oil. For this purpose, solutions of lipid in chloroform were evaporated under a stream of nitrogen, placed in vacuum for 1 h, and oil was added to a final lipid concentration of 0.4 mM. The samples were sonicated for 60 min. Sigma light mineral oil was used to dissolve the lipids for the outer leaflet (emulsion solution) and Roth mineral oil was used as an emulsion oil for the inner leaflet. These specific oils were chosen to optimize vesicle yield and purity. Next, interfacial incubation was carried out by sequentially introducing 250 μL of 700 mOsmol/kg glucose and 200 μL of lipid in Roth oil (interphase lipid-oil solution) in an Eppendorf protein LoBind tube. The tube was left for 3 h and then centrifuged for 5 min at 600×g to ensure the formation of the interfacial lipid layer. To prepare the droplet emulsion, 1 μL sucrose solution (700 mOsmol/kg) was added to 50 μL of lipid in Sigma oil (emulsion solution). Water-in-oil emulsion was formed by performing a series of sequential rubs on a tube rack to form fine lipid droplets. The emulsion was slowly added to the top of the tube with water/oil interface and centrifuged at 130×g for 10 min. The oil layer was pipetted out and the vesicles were collected from the bottom of the tube. All preparation steps were conducted at room temperature (∼24 °C).

To explore control vesicles with symmetric membranes, a standard electroformation protocol was used. Briefly, 10 μl of 4 mM POPC or POPC/POPA mixture with a fluorescent probe (0.2 mol%) in chloroform was spread evenly on indium-tin oxide coated glass coverslips. These were then dried under vacuum for 1 h. The coverslips were then assembled with coated sides opposite to each other and sandwiching a Teflon spacer to form a chamber. The chamber was filled with 2 mL 150 mOsmol/kg sucrose. AC electric field of 1 Vpp and 10 Hz was applied for 2 h, at a temperature of 30 °C. The vesicles were then harvested, and 2-fold diluted in isotonic glucose solution. Osmolalities were measured and adjusted using a freezing-point osmometer (Osmomat 3000, Gonotec, Germany).

Both preparation approaches were evaluated using phosphorus analysis [44] of the measured lipid mass content in the obtained GUVs showed similar lipid amounts in the samples (see Section 8 in SI).

To form large unilamellar vesicles (LUVs) for calorimetry measurements, a thin lipid film was prepared by evaporating a lipid solution in chloroform in a glass vial using a stream of nitrogen. The vial was subsequently kept in vacuum for 2 h to remove any remaining traces of solvent. A volume of 1 mL 150 mOsmol/kg glucose solution was then added and the glass vial vortexed for 1 minute to generate multilamellar vesicles (MLVs). Both the glucose solution and the vial were preheated in an incubator at 50 ×C for 10-15 min prior to mixing. To produce LUVs [45], the MLV suspension was extruded at least 13 times through a 100 nm polycarbonate membrane using a mini-extruder (Avanti Polar Lipids, Alabaster-AL). Extrusion was also performed in the incubator (at 50 ×C) to ensure that the lipids in the solution were in the fluid phase.

### 2.3 Expression, purification and labeling of α-synuclein

To obtain plasmid construct for overexpression of 8×His - tagged murine α-synuclein and these proteins fused with mEGFP (monomeric enhanced green fluorescent protein) in bacterial system, first the coding sequence of N-terminal (Met1 – Lys60) α-synuclein fragment was amplified and subcloned from pET3a aSyn murine to pET28 plasmid containing mEGFP coding sequence by double restriction digest (BamHI and EcoRI enzymes) and ligation according to manufacturer protocols. Then the Restriction Free Cloning method was performed to clone the rest of the α-synuclein sequence into the previously prepared construct and obtain a plasmid encoding the complete α-synuclein sequence. The Restriction Free Cloning procedure was performed according to the instruction described by van den Ent & Löwe [46] using the primers designed in https://www.rf-cloning.org/. Then, to obtain a construct encoding α-synuclein fused with mEGFP protein, site-directed mutagenesis was performed to remove the stop codon located between the sequence encoding α-synuclein and the mEGFP protein. The procedure was performed using the Q5® Site-Directed Mutagenesis Kit according to the manufacturer’s protocol. The web tool NEBaseChanger (https://nebasechanger.neb.com/) was used to design the mutagenic primers. To confirm and validate performed cloning, the obtained DNA constructs were subjected to Sanger sequencing with primers specific to T7 promoter and T7 terminator sequences (Microsynth Seqlab GmbH, Germany). The sequences of the primers used in the cloning procedures are provided in Table S1 of the Supporting Information (SI).

Production of recombinant α-synuclein (αSyn) and α-synuclein-mEGFP (αSyn-mEGFP) protein was performed in *Escherichia coli* NiCo21 (DE3) strain. 200 mL of 50 μg/mL kanamycin supplemented LB Miller broth was inoculated with overnight preculture and incubated at 37 °C at 200 rpm agitation to reach culture optical density (λ = 600 nm) of 0.7. Then the protein overexpression was induced by adding IPTG at a final concentration of 100 or 400 μM for αSyn nad αSyn-mEGFP, respectively. αSyn-mEGFP overexpression was carried out at 18 °C at 200 rpm agitation for 18 h, whereas αSyn overexpression was carried out at 37 °C at 200 rpm agitation for 4 h. Next, cells were harvested by centrifugation (10 000×g, 20 min, 4 °C) and lysed by re-suspending the pellet in 10 mL of a lysis buffer (10 mM HEPES, 500 mM NaCl, 1 mg/mL lysozyme, 0.75% Triton X-100, 25 U/mL OMNI Nuclease, 10 mM imidazole, 1 mM PMSF, 1× Price™ Protease Inhibitor Tablets EDTA free, pH 8.0) and subsequent incubation for 1 h at 4 °C under gentle mixing. Afterwards, the suspensions were sonicated on ice for 15 min at 80 % amplitude and 0.5 cycle (Hielscher UP100H Ultrasonic Processor with MS3 sonotrode). Lysates were then clarified by centrifugation (35 000×g, 30 min, 4 °C) and the supernatants were incubated with 1 mL of previously equilibrated Price™ High Capacity Ni-IMAC resin (αSyn-mEGFP) or TALON® Metal Affinity Resin (αSyn) for 2 h at 4 °C under gentle mixing. For αSyn-mEGFP purification resin was packed into a chromatography column and washed with 100 mL of Wash Buffer 1 (10 mM HEPES, 500 mM NaCl, 10 mM imidazole, 20% glycerol, pH 8.0), 100 mL of Wash Buffer 2 (10 mM HEPES, 300 mM NaCl, 10 mM imidazole, 20% glycerol, pH 8.0) and 100 mL of Wash Buffer 3 (10 mM HEPES, 300 mM NaCl, 10 mM imidazole, pH 8.0) until the absorbance (at λ = 280 nm) of flow through buffer, measured in 1 cm optical path quartz cuvette, decreased below 0.01. Then protein was eluted from resin with an elution buffer (10 mM HEPES, 300 mM NaCl, 200 mM imidazole, pH 8.0). Eluted fractions were then dialyzed against 10 mM HEPES + 150 mM NaCl buffer pH 7.4 and the protein concentration was determined spectrophotometrically (Cary 1E UV-Visible Spectrophotometer) employing excitation coefficient (at λ = 280 nm) calculated using ProtParam tool (web.expasy.org/protparam). For αSyn purification resin, was packed into a chromatography column and washed with 100 mL of Wash Buffer A1 (10 mM HEPES, 500 mM NaCl, 10 mM imidazole, pH 8.0), 200 mL of Wash Buffer 3 (10 mM HEPES, 300 mM NaCl, 10 mM imidazole, pH 8.0) until the absorbance (at λ = 280 nm) of flow through buffer, measured in 1 cm optical path quartz cuvette, decreased below 0.01. Then protein was eluted from resin with an elution buffer (10 mM HEPES, 300 mM NaCl, 200 mM imidazole, pH 8.0). Eluted fractions were then subjected to buffer exchange to PBS pH 7.4 on Econo-Pac® 10DG column. Buffer exchange was performer accordingly to manufacturer protocol. Protein concentration was then determined using BCA assay as described in manufacturer protocol. The purity and molecular weight of the produced protein was determined by SDS-PAGE in Laemmli system [47] (12% and 4-20% gradient resolving gel for αSyn-mEGFP and αSyn respectively) with the following Coomasie Brilliant Blue R-250 staining. The SDS-PAGE analysis is shown in SI Figure S1. The purified protein was than aliquoted, flash frozen in liquid nitrogen and stored at – 80°C.

In addition to αSyn-mEGFP, we employed a second fluorescently labeled analog of the protein, namely αSyn-RED. The labelling was performed using Protein Labeling Kit RED-NHS 2^nd^ Generation according to the manufacturer protocol with minor modifications. Briefly, a solution of 10 μM αSyn in PBS buffer pH 7.4 was incubated for 30 min at room temperature in the presence of 60 μM amine-reactive fluorescent tag RED-NHS. Subsequently, the labelled protein was separated from the unbound fluorophore using size exclusion chromatography column provided by the manufacturer in the kit and equilibrated with 10 mM HEPES buffer pH 7,4 with 150 mM NaClbuffer. Protein concentration and degree of labelling was then analyzed as described in the kit manual.

### 2.4 Fluorescence quantification

The quantification of the fluorescence signal of the NBD-PC dye was done on a confocal microscope (Leica microsystems TCS SP5, Wetzlar, Germany) using a 40× HCX PLAN APO dry objective, NA 0.75. Confocal cross-section images of 512 px × 512 px were collected. Identical acquisition settings (constant zoom, laser intensity, detector gain) were maintained for all measurements. NBD-PC and Atto488-DOPE were excited with an argon laser line at 488 nm (10 % laser intensity) and the emission signal was collected in the range 500–600 nm. TexasRed-DHPE, Rh-DOPE, Rh-DHPE and DilC_18_ were excited at 550 nm and emission signal was collected in the range 565-730 nm. The quantification of αSyn-mEGFP and αSyn-RED fluorescence was performed on an TCS SP8 confocal microscope (Leica microsystems, Wetzar, Germany) using a 63× oil immersion objective, NA 1.40, and on a Stellaris confocal microscope (Leica microsystems) using a 86× water immersion objective, NA 1.20. Samples with αSyn-mEGFP were excited with a 488 diode laser and the emission signal was collected in the range 500–600 nm; samples with αSyn-RED were excited at 640 nm and the emission collected in the range 650-755 nm. The vesicles were 2-fold diluted in isotonic solution of glucose. Both αSyn-mEGFP and αSyn-RED (stored in 10 mM HEPES 7.4 pH with 150 mM NaCl) were diluted 1:1 with isotonic glucose and added to a final concentration of 0.5 μM (typically 3-5 μl of protein solution was added to 300 μl vesicle suspension), followed by incubation for 5 min to ensure homogeneous distribution of the protein within the sample. The final protein concentration was chosen based on optimization tests on electroformed GUVs containing various molar fractions of PA (see SI Figures S2 and S3). The average fluorescence intensity over the whole contour of the GUV (with thickness of 5 pixels) was evaluated using the Circle Skinner plugin of ImageJ (github.com/tinevez/CircleSkinner). The average pixel intensity in the vesicle interior was manually measured and the value subtracted from that of the membrane contour. The measurements were performed at 23±1°C.

### 2.5 Zeta-potential measurements

Zeta potential measurements on GUVs were done according to already established protocols [48, 49] using Malvern ZetaSizer NanoZS (Malvern, UK) and disposable folded capillary cells (DTS1070; Malvern Panalytical). Briefly, GUVs were measured in both dip-cell (Malvern ZEN1002, with the voltage for electrophoretic movement set to 10 V) and U-cells (Malvern DTS1070, with voltage set to 150 V). The GUV samples were measured not more than 3 times as repetition can result in GUV rupture at the electrodes. Measurements showing poor quality report by the Malvern software were discarded. At least three independently obtained populations were measured to ensure reproducibility of the reported values. Sucrose/glucose solutions were supplemented with 5 mM NaCl; measurements at different salinity are provided in the SI (Figure S4).

### 2.6 Micropipette aspiration

Micropipettes with inner diameter of 5–10 μm were prepared from glass capillaries (World Precision Instruments, Sarasota, FL, USA) using of a micropipette puller (Sutter Instruments USA, Novato, CA) and their tips were shaped with a microforge (Narishige, Tokyo, Japan). Before use, each micropipette was coated with 1 mg/mL casein solution to prevent vesicle adhesion to the glass. To apply suction pressure, the micropipette was connected to a water reservoir mounted on a vertical translational stage (M-531.DD; PI, Karlsruhe, Germany). Manipulation in the sample was achieved with the use of micromanipulators (MHW-103 or MLW-3; Narishige, Japan) secured to coarse manipulators (MMN-1; Narishige). Vesicles were visualized on Leica SP5 (Leica, Germany) confocal microscope. 1024 px × 1024 px images were collected using a HCK PLAN APO 40× NA 0.75 dry objective. The TexasRed-DHPE dye in the membrane was excited at 594 nm and emission collected in the range 600-700nm. To avoid concentration changes resulting from evaporation during longer observation times, the chamber opening was covered with a layer of oil (Sigma light mineral oil). Upon aspiration, the vesicle was left to equilibrate for 3 min before changing the suction pressure. The size of the vesicle spherical cap outside the pipette and the length of aspirated part was measured from the images using custom-written script in MATLAB (The MathWorks, Natick, MA), which automatically selects a spherical cap portion and calculates the vesicle radius in each frame. The script is designed in such a way that for each frame the region covering at least half of the vesicle is manually selected. This is followed by Taubin nonlinear circle fitting to obtain the vesicle radius for 1%, 2% and 3% pixels of highest intensity value. The average radius is taken further into calculations to obtain the area compressibility. The error in tension and area change were estimated from the errors of the individual input parameters and error analysis based on differential approach. The experiments were performed at room temperature (23±1 °C).

### 2.7 Molecular dynamics simulations

The full-atomistic MD simulation was performed using NAMD 2.13 [50] software with CHARMM36 force fields [51, 52] under NPT conditions (constant number of particles, pressure, and temperature). Membrane systems were prepared from 648 lipid molecules (324 lipids per leaflet). Octane was parametrized using the CGENFF force field [53] and inserted into pre-equilibrated membranes. The systems were hydrated with 75 water molecules per lipid molecule and the charged lipids were neutralized with positive counter ions. A standard equilibration procedure was used [54]. The total simulation time was at least 200 ns, of which the last 10 ns were used for analysis. Simulations were carried out at ∼22 °C (295 K). The membrane thickness h_pp_ was calculated as the difference between the mean height values of phosphorus atoms in opposite leaflets. Density profiles were plotted with VMD density profile tool [55]. The stretching elasticity modulus *K*_*A*_ was calculated with a method developed by Doktorova *et al*. [56]. Interdigitation was calculated with MEMBPLUGIN [57] in which it is defined as the width of the overlap of two leaflets mass distributions along the membrane normal.

### 2.8 Differential scanning calorimetry (DSC)

The thermal profiles of LUV solutions were measured using a VP-DSC scanning calorimeter (MicroCal, Northampton, MA). The reference cell was filled with approximately 0.5 mL 150 mOsmol/kg glucose and the sample cell was filled with approximately 0.5 mL solution of LUVs composed of POPC:POPA 80:20 (5 mM total lipid concentration) or DPPC (10 mM lipid concentration).. The heating rate was set to 20 °C/h. Baseline subtraction was performed in Microcal Origin 7.0.

### 2.9 Statistics

To determine whether there was a significant difference between the parameters, the Kruskal-Wallis ANOVA test was used, with a significance level of 0.05 unless otherwise specified. Non-parametric post-hoc tests were conducted using the npposthoc add-on in OriginPro 2015 (OriginLabs) software. Weighted average values were calculated, taking into account the measurement error, and were presented alongside the weighted standard deviation. The weight assigned to each measurement was determined by calculating its inverse error.

## 3 RESULTS AND DISCUSSION

### 3.1. Preparing asymmetric POPA vesicles and verifying their membrane asymmetry and composition

We employed the inverted emulsion method [41] to prepare asymmetric GUVs from POPC and POPA lipids. Using an adapted approach with modifications described in the experimental section, we obtained high yield of vesicles without visible defects. Concerns have been raised in the literature about the final lipid composition of vesicles obtained with the inverted emulsion method and the presence of residual oil, which could potentially impact the properties of the obtained membranes [58, 59]. To address these concerns, we employed several techniques to characterize the prepared vesicles and probe their membrane composition.

First, we performed fluorescence intensity measurements to investigate whether the vesicles exhibited asymmetry. We incorporated the fluorescent probe NBD-PC to the leaflet containing POPA and compared the fluorescence intensity of both symmetric and asymmetric vesicles. The asymmetric membranes are expected to exhibit half the intensity value of the symmetric ones. Indeed, as shown in Figure 1A, we observed an intensity drop roughly by half in the asymmetric system, suggesting the presence of leaflet composition asymmetry. As shown in Figure 1A the fluorescence intensity drops by roughly half in the asymmetric system. If all lipids were accordingly incorporated in the intended leaflet, this observation indicates indeed an asymmetric membrane composition. However, the observed reduction in membrane intensity might be associated with the twice lower fraction of dye used in the case of the asymmetric membranes. To further probe for the asymmetric character of the membranes we employ protein the αSyn-mEGFP.

**Figure 1.**
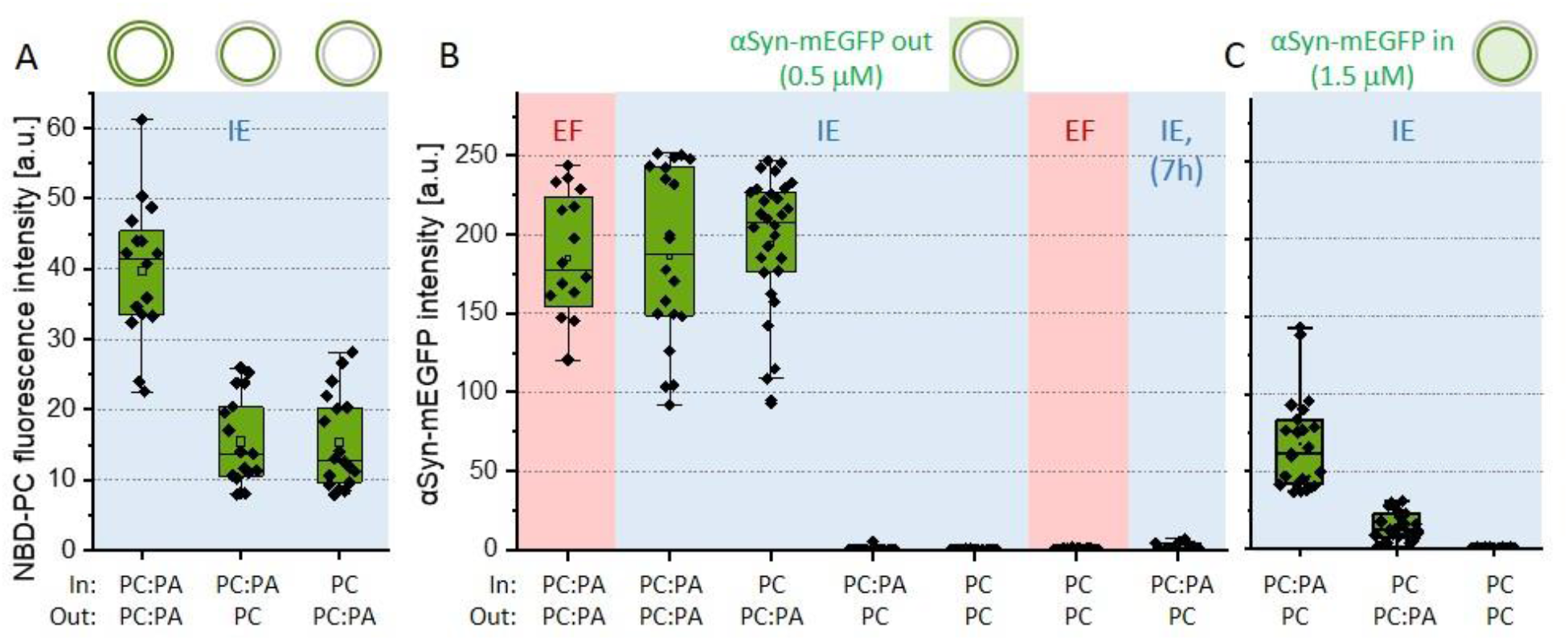
Assessing the membrane asymmetry from the fluorescence signal of incorporated NBD-PC and soluble αSyn-mEGFP. Internal and external leaflets (labelled as “In” and “Out”) with composition POPC:POPA 80:20 (molar ratio) are indicated as PC:PA, and pure POPC as PC. (A) NBD-PC fluorescence intensity of symmetric and asymmetric membranes in which the PC:PA leaflets are labelled with 0.5 mol% NBD-PC (green) as schematically indicated above the graph; all samples were prepared with the inverted emulsion (IE) method (highlighted in light blue). (B) Fluorescence signal of 0.5 μM αSyn-mEGFP added externally to symmetric and asymmetric vesicles prepared with electroformation (EF, pink background) or the inverted emulsion method (IE, light blue). The last set of data shows the intensity after incubating the vesicles with the protein for 7 h. The protein intensity on the membrane shows signal from PA in the external vesicle leaflet as schematically illustrated above the graph. (C) αSyn-mEGFP fluorescence measured on vesicles, which were prepared to encapsulate the protein at concentration of 1.5 μM. The protein intensity on the membrane shows signal from PA in the internal vesicle leaflet as schematically illustrated above the graph. All measurements were performed at room temperature (24 °C). Boxes heights show lower and upper quartile (25-75%), line in box shows median value and square point - average value. Bars represent upper and lower whisker (1.5 IQR value) and diamond points represent measurements on individual vesicles.

To further characterize the leaflet compositions and in particular the distribution of POPA, we used the protein αSyn-mEGFP, which is known to bind with high affinity to membranes with PA and act as a PA sensor [39, 40]. Figure 1B shows the results of adding the protein (to a final concentration of 0.5 μM) to solutions of asymmetric vesicles obtained with the inverted emulsion method (see SI Section 2 and Figure S2 and S3 for optimization steps); as a control, we examined (symmetric) electroformed vesicles. In the case where POPA was in the external leaflet, the vesicles exhibited high fluorescence signal of value comparable to that of the control. Hardly any signal was detected when the POPA-containing leaflet was the internal one. We also confirmed the stability of the membrane system by measuring vesicles with POPA on the inner leaflet 7 h after incubation (Figure 1B). The lack of significant signal increase suggests that there is neither substantial flip-flop of POPA from the inner to the outer leaflet nor translocation of αSyn-mEGFP to the vesicle interior during this time.

To probe the internal leaflet of the vesicles, we added 0.5 μM αSyn-mEGFP to the aqueous phase of the water-in-oil emulsion forming the vesicle interior. The protein signal at the membrane surface was found very low compared to that of vesicles with externally added protein (Figure 1C). This result could imply that the amount of POPA incorporated in the inner leaflet is much smaller than the one we could incorporate in the outer one, but this hypothesis is questionable considering the results for NBD-PC in Figure 1A. We speculated that the low signal compared to that when probing POPA in the outer leaflet is associated with the larger external pool of protein available for binding in the latter case; we also cannot exclude partial protein damage or loss during the emulsification step for the water-in-oil emulsion forming the inner leaflet. To this end, we increased the concentration of αSyn-mEGFP inside the vesicles threefold. This implies that these measurements (Figure 1C) cannot be quantitatively compared to data obtained with externally added protein (Figure 1B). We observed a significant intensity drop when removing POPA from the inner leaflet. Interestingly, the fluorescence value for the system with POPA targeted to the outer leaflet was not negligible. Since we could exclude significant POPA flip-flop as well as protein translocating across the membrane as suggested by the long-time observations in Figure 1B (see also Figure S4C), we presume that a fraction of POPA relocates to the inner leaflet already during the preparation step of emulsion transfer. However, judging from the data obtained on electroformed and emulsion-transfer symmetric vesicles in Figure 1B and the nonlinear fluorescence dependence on PA concentration (Figure S2B), this fraction of relocated POPA is not substantial. Overall, the data obtained with αSyn-mEGFP, cannot be used to unequivocally confirm the precise amount of POPA in the inner membrane leaflet.

To validate the conclusions drawn from our fluorescence-based experiments and to characterise the systems further, we explored the surface charge of the vesicles using zeta-potential measurements as an independent method. This technique is not commonly used with GUVs due to uncertainties such as the unknown history of the GUVs with respect to prior rupture and leaflet mixing, potential collapse at the electrodes, etc. However, it has been successfully used to assess the zeta-potential of giant vesicles in previous studies [48, 49]. To avoid electrode polarization and field distortion, in these measurements, the GUV suspensions were supplemented with NaCl (final concentration of 5 mM). The data are presented in Figure 2; further details and results for vesicles in the absence of salt and in the protein buffer are given in SI Section 4. The symmetric POPC vesicles obtained via electroformation and emulsion transfer, as well as the asymmetric ones with POPA in the inner leaflet and only POPC on the outer one, yield similar results for the zeta potential (the values indicate relatively high negative surface charge but are consistent with data reported by Carvalho et al. [49]). They also suggest negligible transfer of POPA from the inner to the outer leaflet. Similarly, symmetric and asymmetric GUVs prepared with both methods but containing POPC:POPA 80:20 in the outer leaflet exhibited similar and more negative zeta-potential values. We probed whether the zeta potential of the asymmetric vesicles would change over time as a result of inter-leaflet exchange. No substantial differences were detected after 4 h (Figure S4C). This corroborates our conclusion for absence of significant PA flip-flop during this time. Overall, the results support the conclusion drawn from fluorescence quantification of the membrane asymmetry and suggest that the investigated systems correspond relatively well to the intended composition of the outer leaflet. With this established, we now proceed with investigating the mechanical properties, phase state and lateral organization in the membranes.

**Figure 2.**
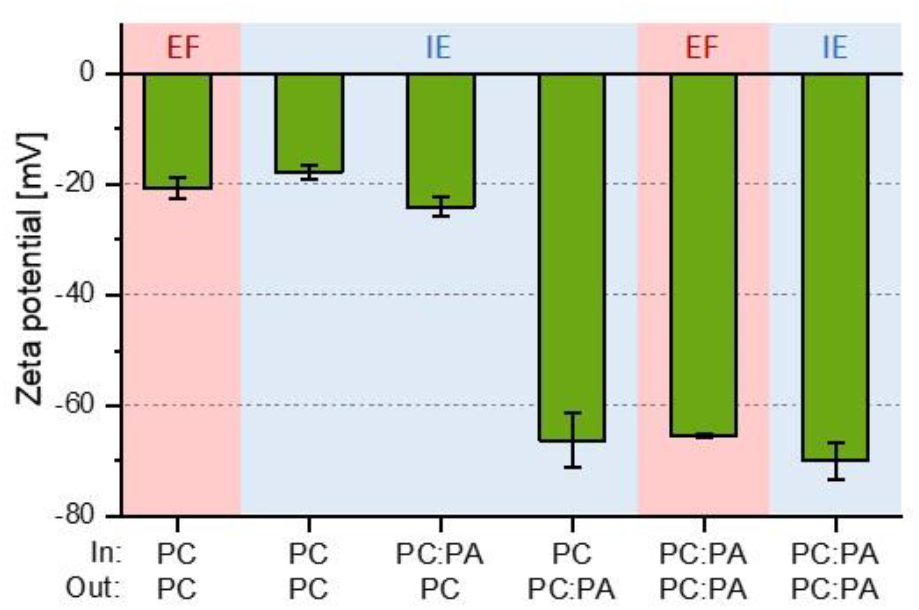
Zeta potential of GUVs prepared via electroformation (EF, highlighted in pink) and inverted emulsion (IE, light blue) methods. Internal and external leaflets (In and Out) with composition POPC:POPA 80:20 (molar ratio) are indicated as PC:PA and pure POPC as PC. The external solution was sucrose/glucose with 5 mM NaCl. The measurements were performed at 25°C.

### 3.2. Mechanical properties of symmetric vs asymmetric membranes

We measured the membrane area compressibility moduli, *K*_*A*_, using micropipette aspiration of GUVs doped with a small fraction of TexasRed DHPE (Figure 3A,B). The vesicles were pre-stressed to ensure no contributions from area stored in nanostructures [60]. Hysteresis tests were also performed confirming that the membrane does not adhere to the pipette and no effects with long-time observation (e.g. evaporation from the chamber) are present, see SI Section 5 and Figure S5.

**Figure 3.**
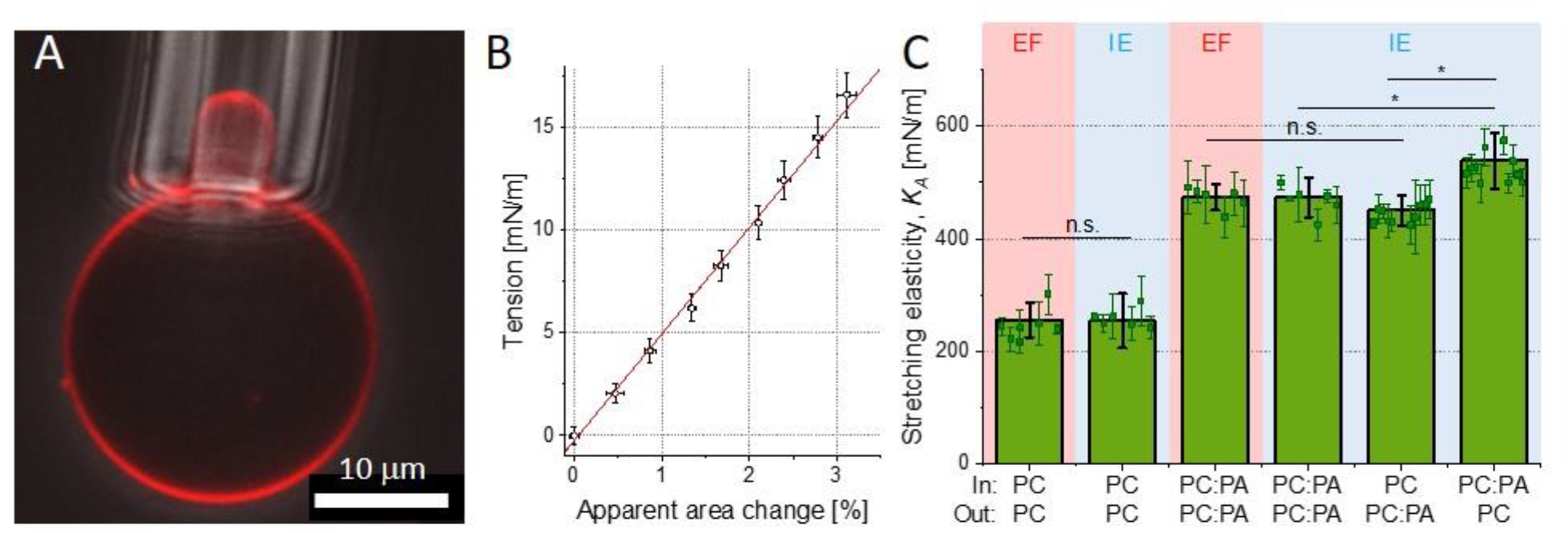
Micropipette aspiration measurements of the stretching elasticity modulus *K*_*A*_. (A) Snapshot of an asymmetric GUV (with POPC:POPA 80:20 in the inner leaflet, POPC in the outer leaflet and 0.2 mol% TexasRed-DHPE in both leaflets) aspirated in a micropipette: overlay of a confocal cross section (showing the fluorescently labelled membrane) and a phase contrast image (showing the micropipette tip). (B) Tension-area expansion plot for the vesicle shown in (A). The stretching elasticity modulus obtained from the slope of the data is (500±25) mN/m. (C) Area compressibility moduli determined for all investigated vesicle systems prepared via electroformation (EF, pink background) and inverted emulsion (IE, light blue) methods. Internal and external leaflets (In and Out) with intended composition POPC:POPA 80:20 (molar ratio) are indicated as PC:PA and pure POPC as PC; note that while the POPA fraction in the external leaflet corresponds well to the intended one, the precise POPA fraction of the internal leaflet of IE GUVs is unclear. At least 10 vesicles were measured for symmetric and 15 for asymmetric membranes; the individual measurements are shown with symbols and standard deviations. The green bars show mean values and standard deviations over the populations. All measurements were performed at room temperature (23±1°C). Statistical significance was determined with Kruskal-Wallis ANOVA test and followed by a post-hoc test. * represents a significant difference across membrane compositions (p<0.05), while lack of a significant difference (p > 0.05) is denoted with n.s.

For POPC symmetric vesicles, we found the stretching elasticity modulus to be around 250 mN/m both for electroformed and inverted-emulsion vesicles (Figure 3C). The proximity of these values suggests that potential presence of oil does not detectably affect this membrane mechanical property. The values are consistent with previous reports [19, 61, 62]. For POPC:POPA 80:20 symmetric vesicles, we measured higher area compressibility around 480 mN/m, see Table 1. These data are also consistent with simulations [19]. Importantly, we again observed no statistically significant difference in the stretching elasticity modulus of vesicles prepared via electroformation compared to those prepared via the inverted-emulsion method. This is in agreement with a recent study on another mechanical parameter, the bending rigidity, which was shown not to differ for POPC vesicles prepared with the two methods using similar sugar concentrations [63]. Note that the electroformed vesicles in our study are in sugar solutions of a much lower osmolality (150 mOsmol/kg) compared to vesicles obtained with the phase-transfer method (700 mOsmol/kg). The similar *K*_*A*_ values suggest that in this concentration range, sugars do not detectably affect the stretching elasticity modulus.

**Table 1.** Stretching elasticity modulus, *K*_*A*_, values determined experimentally with micropipette aspiration of vesicles with intended leaflet compositions POPC (PC) and POPC:POPA 80:20 (PC:PA) produced with electroformation (EF) and inverted emulsion (IE) method as shown in Figure 3C, and from molecular dynamics simulations of symmetric membranes (data obtained at 22-24°C). The MD simulation value for symmetric POPC:POPA 80:20 membranes indicated with an asterisk is collected at 30°C and was reported in reference [19].

| Experiment (micropipette aspiration) |  |  |  | Simulations (molecular dynamics) |  |  |
| --- | --- | --- | --- | --- | --- | --- |
| Method | Inner leaflet | Outer leaflet | $K_A$ [mN/m] | Both leaflets | $K_A$ [mN/m] | Thickness, [mN/m] |
| EF | PC | PC | $255 \pm 31$ | PC | $252 \pm 38$ | $3.70 \pm 0.04$ |
| IE | PC | PC | $249 \pm 35$ | PC + octane | $258 \pm 33$ | $3.83 \pm 0.03$ |
| EF | PC:PA | PC:PA | $487 \pm 35$ | PC:PA | $455 \pm 24^*$ | $3.92 \pm 0.03$ |
| IE | PC:PA | PC:PA | $480 \pm 41$ | PC:PA+ octane | $448 \pm 32$ | $4.03 \pm 0.02$ |
| IE | PC:PA | PC | $539 \pm 49$ | | | |
| IE | PC | PC:PA | $451 \pm 35$ | | | |

Interesting if not unexpected results were obtained when comparing the compressibility moduli of symmetric and asymmetric bilayers. Notably, the *K*_*A*_ value we measured for the asymmetric system with POPC:POPA present only in the external leaflet is not significantly different from that for symmetric POPC:POPA membranes. However, the stretching elasticity modulus of the inversed asymmetric system with a POPC:POPA leaflet on the inside, is higher than the values obtained for both the symmetric POPC:POPA membrane and membranes with POPA only in the outer leaflet. Despite being small, the difference is statistically significant. These findings are summarized in Figure 3C and Table 1. Data for individual vesicles are given in Figure S6.

### 3.3. Molecular dynamic simulations of symmetric, octane containing membranes

To validate our experimental findings and understand the potential impact of residual mineral oil on membrane elasticity, we conducted molecular dynamics (MD) simulations on symmetric membranes containing octane. The simulated oil-free membranes were expected to model those of electroformed GUVs, while bilayers containing the oil could mimic the behaviour of inverted emulsion GUV membranes. We did not attempt to establish asymmetric systems for assessing the membrane mechanical properties, as it would require setting up an optimal lipid organization of the asymmetric membrane. Various approaches, such as those based on individual area per lipid, leaflet surface area, or zero leaflet tension (differential stress), have been proposed for such systems (see e.g., [64, 65]). The latter approach [66, 67], while more suitable, would involve laborious iterative adaptations, which is beyond the scope of this work. Therefore, we focused solely on symmetric bilayers to investigate the specific role of oil impurities. Provided the simulations show reasonable agreement with the experimental data on stretching elasticity, we were hoping to employ them to explore further membrane properties inaccessible to experiments.

Mineral oils used in phase-transfer methods for GUV preparation are mostly composed of n-alkanes. Therefore, following an established protocol [68], we modelled 8-alkane (octane) molecules and inserted them into the acyl chain region of an equilibrated membrane. As there are no clear indications of the amount of oil residue in lipid membranes prepared with emulsion transfer, we used a generous oil fraction of 100 octane molecules and 653 lipid molecules corresponding to ∼13 mol% of oil. This choice is likely a significant overestimation but was meant to establish a clear trend.

Figure 4 displays snapshots of the four systems along with their density profiles. In the octane-free membrane (Figure 4A, B, E, F), the density profiles of the acyl chain region reveal two maxima. In the presence of octane (Figure 4C, D, G, H), these maxima become more pronounced, suggesting that the oil residues affect acyl chain organization, thereby influencing the positions of lipids within the membrane.

**Figure 4.**
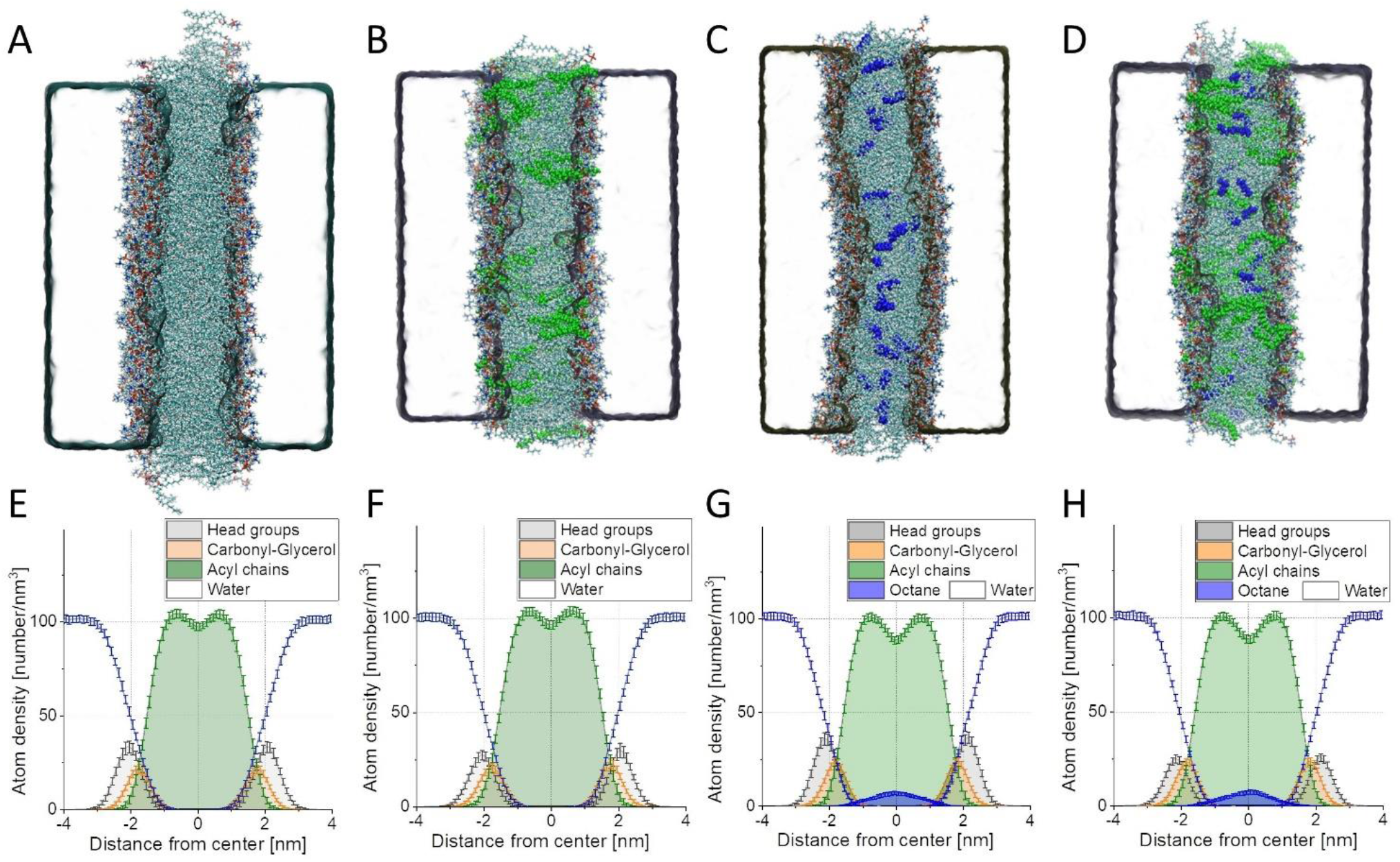
MD simulations of symmetric POPC and POPC:POPA 80:20 planar membranes without (A, B) and with ∼13 mol% octane (C, D). (A-D) Snapshots of the bilayers: The POPC molecules are shown colored according to atom type, POPA is shown in green and octane in blue. POPC is colored by atom type. (E-H) Density profiles of the investigated asymmetric systems.

We then assessed the membrane elasticity of the simulated bilayers. For this, we employed a method developed by Doktorova et al. [56]. It has the advantage of analysing *K*_*A*_ based on matching the leaflet area per lipid, thus allowing to determine *K*_*A*_ separately for each leaflet. While this approach is still debated [69], it appears to be most suitable given the simulated systems. Results of determined compressibility are displayed in Figure 5 and comparison with experimental data are presented in Table 1.

**Figure 5.**
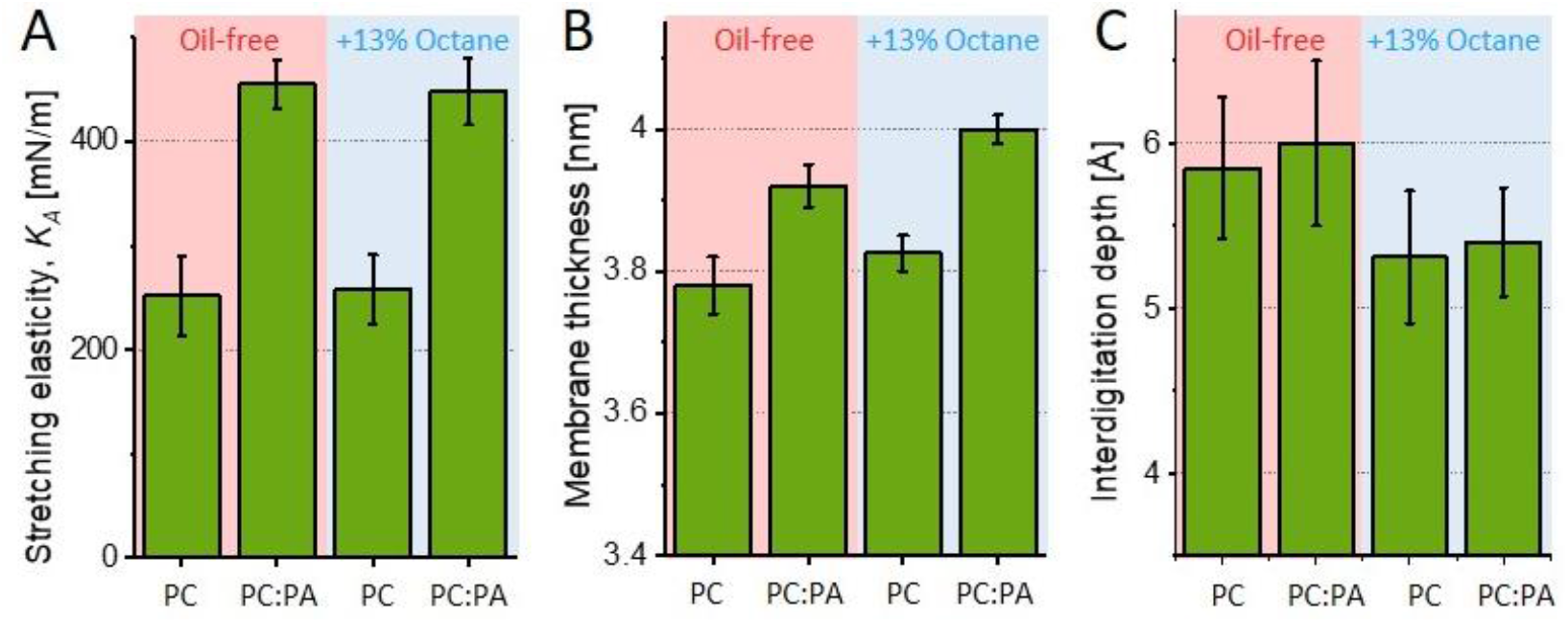
MD simulation values for the stretching elasticity, membrane thickness and interdigitation depth for symmetric POPC and POPC:POPA 80:20 membranes, which are either oil-free (pink background) or containing ∼13 mol% octane (light blue background) mimicking the respective conditions of electroformed and inverted-emulsion preparation of GUVs. (A) Stretching elasticity modulus *K*_*A*_. No significant difference between the systems with and without octane are observed. (B) Membrane thickness values. Presence of octane results in increased thickness. (C) Calculated interdigitation depth shows reduced interleaflet coupling in the presence of octane. The data for the oil-free POPC:POPA 80:20 system were taken from [19].

Several observations, arising from the comparison between experiments and simulations, are worth emphasizing. First, symmetric POPC:POPA 80:20 bilayers have higher elasticity modulus compared to that of pure POPC membranes (consistent with previous reports [19, 62]) and the absolute values obtained with our experiments and the simulations are in agreement within the range of uncertainty. This not only implies that simulations correctly represent the symmetric experimental system but also that they support the hypothesis of specific POPA incorporation in the membrane leaflets. Second, the presence of octane up to an oil-to-lipid ratio of ∼13% did not significantly alter the membrane compressibility, consistent with the experimental data for symmetric membranes prepared with via electroformation and inverted emulsion methods (Table 1). Third, as could have been expected, membrane thickness increased due to the presence of oil residues, as shown in Figure 5B. This octane-induced thickness increase is associated with reduced leaflet coupling as visualized from the decreased interdigitation depth as shown in Figure 5C (the thickness increase corresponds to twice the decrease in interdigitation depth). This finding sparks curiosity, because together with the unaltered stretching elasticity modulus, the increased thickness suggests that oil residues could affect, namely increase, the bending rigidity of membranes produced with the inverted emulsion method; note that the stretching and bending elasticity moduli *K*_*A*_ and κ are related to the membrane thickness *h* as κ/*K*_*A*_ ∼ *h*^2^/α, where α is a constant reflecting the interleaflet coupling [70-72]. Strictly speaking, we cannot rule out the possibility that in our GUV-based systems the fraction of oil retained in the membrane depends on the specific lipid composition chosen. Exploring this issue would necessitate precise compositional analysis, which is beyond the aims of our current study. The results, however, emphasize that experimental methods to form asymmetric lipid membranes and data obtained from them should be treated with extreme caution.

### 3.4. Phase separation in POPC:POPA membranes induced by DOPE-based fluorescence dyes

The main phase transition temperature of pure POPA membranes is 28°C [73] and that of POPC membranes is -2°C. When mixed with POPC at 1:1 ratio, the phase transition of POPA is suppressed in the range 10-70°C, unless calcium ions are present [74]. Our own differential scanning calorimetry measurements of POPC:POPA 80:20 membranes also did not show a phase transition in the range between 10 and 50°C (Figure S7). This is consistent with the MD simulations, which showed lack of lipid clustering into domains. Thus, at room temperature, which is the temperature of our microscopy observations, no phase separation in the membranes is expected for the binary mixture. Indeed, confocal microscopy inspection of 3D scans of the vesicles labeled with TexasRed-DHPE showed homogeneous membrane (Figure 6A).

**Figure 6.**
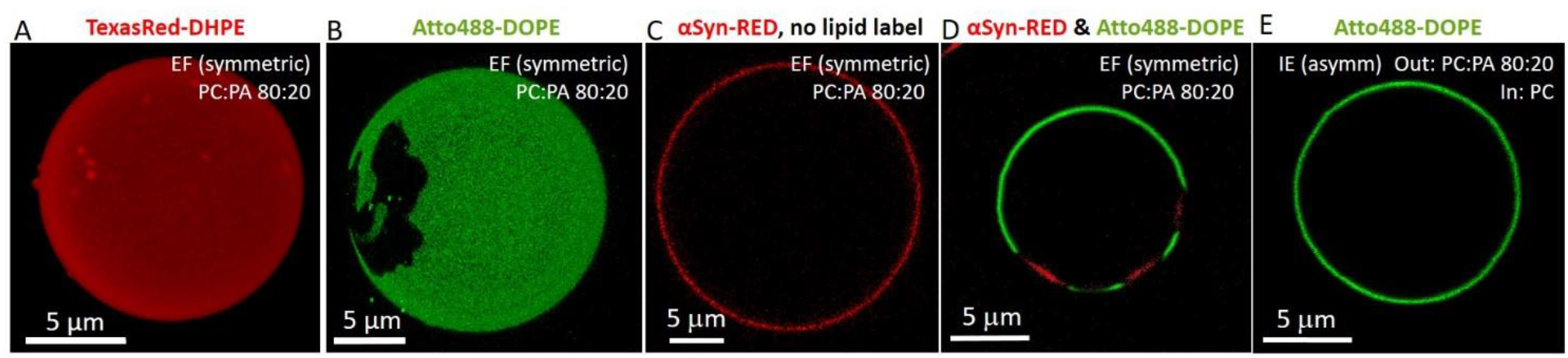
Inclusion of DOPE-based dye (at 0.5 mol%) in homogeneous POPC:POPA 80:20 membranes induces domain formation, which is suppressed in asymmetric membranes containing the dye only in either one or both leaflets. (A) 3D confocal projection image of electroformed POPC:POPA 80:20 vesicle labeled with 0.5 mol % TexasRed-DHPE (false red) exhibits homogeneous membrane. (B) Upon inclusion of 0.5 mol% Atto488-DOPE (green), dark gel-like domains are observed. (C) No phase separation in the label-free membrane is observed, as shown by the homogeneous distribution of externally added 0.5 μM αSyn-RED (red). (D) When αSyn-RED is added to phase-separated GUVs (like the one shown in panel B), the protein PA sensor binds predominantly to the Atto488-DOPE-depleted domain indicating that it is enriched in POPA. (E) Inverted emulsion vesicle with asymmetric leaflet composition with outer leaflet POPC, and inner leaflet POPC:POPA 80:20 labeled with Atto488-DOPE, exhibits homogeneous membrane, contrary to symmetric POPC:POPA vesicles as exemplified in panel (A).

Since the use of fluorescent labels for vesicle visualization and characterization as well as phase state description is abundantly used in GUV literature, we explored other widely used fluorescent markers. Tests with NBD-PC, Rh-DHPE and DilC_18_ also did not show any domain formation in the membrane (see Figure S8A-C, for clarity we give confocal cross sections even though the whole vesicle surface was examined). Surprisingly, when a small fraction (0.5 mol %) of Atto488-DOPE or Rh-DOPE was incorporated into the membrane, confocal microscopy observations revealed the presence of dark domains in symmetric POPC:POPA 80:20 vesicles prepared using both electroformation and emulsion transfer (examples shown in Figure 6B and Figure S8D-E). The irregular shape and stability of these domains suggest gel-like nature. They exhibited no shape change over time and showed lateral displacement, indicative of the surrounding phase being fluid. Similar dark domains were observed for lower fractions of POPA, namely POPC:POPA 95:5 and 90:10 (as exemplified in Figure S8F), albeit with a slightly smaller area. Our investigation revealed that only two fluorescent probes, namely Atto488-DOPE and Rh-DOPE, both DOPE-based, resulted in gel-like domain formation, contrary to membranes labeled with TexasRed-DHPE, NBD-PC, Rh-DHPE and DilC_18_ (Figure 6A and Figure S8A-C).

These observations raise the question about the actual phase state of the dye-free GUV membrane (note that the DSC measurements showing no phase transitions were conducted on LUVs, Figure S7). To eliminate the effect of fluorescent lipid probes on the membrane phase state and appearance of domains, we prepared dye-free GUVs and incubated them with αSyn labelled with Red-NHS (αSyn-RED, 0.5 μM final concentration). The PA-sensor protein was found to bind homogeneously on the POPC:POPA vesicles, indicating lack of phase separation (Figure 6C). This is consistent with simulations and calorimetry data and indicates that the dyes correctly represent the phase state of the membrane and do not show domains, contrary to DOPE-based labels. Upon protein incubation of phase-separated POPC:POPA 80:20 vesicles labeled with Atto488-DOPE, αSyn-RED marked the dark gel-like domains on the membrane (Figure 6D). This indicates that the gel-domains are PA-rich, which is understandable considering the higher transition temperature of POPA.

PE lipids have been reported to strongly interact with PA via hydrogen bonding between the primary amine in the head group of PE and the phosphomonoester head group of PA enhancing PA deprotonation and increasing its negative charge [75, 76]. We thus explored whether addition of a small fraction (0.5 mol%) of DOPE to GUVs (POPC:POPA:DOPE 79.5:20:0.5) labelled with 0.5 mol% TexasRed-DHPE (the dye that previously showed homogeneous distribution, lack of domains) will result in phase separation. Indeed, the addition of this small fraction of DOPE resulted in phase separated vesicles (Figure S8G). Presumably, DHPE-bonded to TexasRed (as in TexasRed-DHPE) behaves less similar to DOPE compared to Atto488-DOPE and Rh-DOPE, most probably as a result of differences in heagroup-substituting fluorophores and/or acyl chain configuration.

We also explored the persistence of the gel domains (induced by the DOPE-based dyes) in asymmetric membranes prepared with the inverted emulsion method. TexasRed-DHPE was added to both leaflets, to outer leaflet and to inner leaflet (Figure S8H-I). No domains were observed irrespective of the leaflet compositional order - POPC:POPA 80:20 in the internal or external leaflet and POPC in the external or internal respectively or the location of the DOPE dye (Figure 6E, Figure S8H-I). This observation is consistent with a recent report demonstrating that in asymmetric bilayers, the liquid-disordered leaflet dominates the phase state of the whole membrane [77]. This report together with our results, emphasize the presence of leaflet coupling in asymmetric membranes, also shown in a number of studies on cholesterol containing membranes [78, 79].

## 4 DISCUSSION

Several experimental approaches were employed to characterize the composition of the leaflets of symmetric and asymmetric membranes. Based on fluorescence quantification using αSyn-mEGFP and on zeta potential measurements (Figures 1 and 2), we found that the composition of the outer leaflet in vesicles formed by both electroformation and inverted emulsion methods is identical within the measurement accuracy. Long-term observations of the asymmetric vesicles (Figure 1B and Figure S4C) suggest no detectable transfer of POPA from the outer leaflet to the inner one after GUV formation has been completed.

The composition of the inner leaflet is challenging to characterize due to methods limitations. Compared to the outer leaflet, the signal of encapsulated αSyn-mEGFP binding to the inner leaflet is low even after a threefold increase in protein concentration (see Figure 1C). This observation could be attributed not only to a potential difference in the POPA fraction from the intended value, but also to variations in lipid-to-protein concentration conditions, potential damage, aggregation and/or loss of the protein during emulsification, or, given the sensitivity of POPA to pH, alterations in surface charge of the leaflet and hence modified affinity of αSyn towards PA. Consequently, we can only infer the presence of POPA in the inner leaflet, without the ability to quantify the exact amount or specify whether it is lower, higher, or the same as in the target composition.

The stretching elasticity *K*_*A*_ measured experimentally with micropipette aspiration of GUVs featuring symmetric POPC and POPC:POPA membranes is consistent across different GUV preparation methods (Figure 3 and Table 1). This suggests that, if oil residues are present, they do not substantially alter the membrane elasticity. This finding was further validated through simulations of symmetric bilayers containing octane as an oil-residue representative (Figure 4). The experimental *K*_*A*_ value for symmetric POPC:POPA 80:20 membranes is significantly higher than that of POPC membranes (for both GUV preparation approaches), consistent with simulations of both oil-free and oil-doped bilayers (see Table 1). This suggests that, first, the simulation settings are appropriate for predicting the elasticity of bilayers with charged PA lipids at different fractions, and second, that oil impurities at a fraction as high as 13 mol % do not affect the stretching elasticity of the membrane. This outcome also indicates that stretching elasticity measurements are not a reliable indicator for detecting the presence of such impurities in the membrane.

The simulations (Figure 5) also showed that while the stretching elasticity is not affected by the presence of oil, the membrane thickness and interleaflet coupling (evidenced by the interdigitation depth) are. The larger thickness of the oil-doped membranes might imply higher bending rigidity of the membranes produced using the inverted emulsion method. Of course, the degree of this outcome could depend on the type of lipids and oils used. The reduced leaflet coupling in the presence of oil (Figure 5C) might be related to observations showing that residual oil in GUVs prepared with the inverted emulsion method destabilize porated membranes [38].

The stretching elasticity was also measured on GUVs with asymmetric membranes. Membranes with POPC:POPA in the outer leaflet showed similar elasticity to symmetric POPC:POPA membranes. However, based on the presented data we cannot exclude that the inner leaflet of the asymmetric membrane might have been populated with POPA suppressing the asymmetry.

This reasoning gains support when considering the individual steps of the inverted emulsion preparation protocol. It is important to emphasize that the incorporation of individual lipids into the targeted leaflet can occur in a similar manner only if they have the same affinity to the water-oil interface and similar or sufficient time to establish this partitioning from the oil to the interface. However, the formation of the two leaflets in the asymmetric GUVs follows different pathways. In particular, they are allowed different amounts of time to equilibrate when prepared with the inverted emulsion method. In addition, the monolayers are also formed under conditions of different ratio between the volume of the oil phase and the area of the water-oil interface to which the lipids (that might coexist as free species and inverted micelles of different mobility) have to relocate and saturate. In the case of the inner leaflet, the water-in-oil emulsion is prepared and used within a few seconds (to avoid droplet coalescence) and is characterized by a much larger area of the water-oil interface (i.e. the combined surface area of all the emulsion droplets) compared to the situation of the outer leaflet. The outer leaflet, on the other hand, is assembled from a well equilibrated (over a few hours) oil-water interface of smaller area (roughly the cross-section of the centrifugation tube) and a larger bulk phase.

Considering the differences in leaflet formation conditions mentioned above, we conclude that free lipids in the oil phase that forms the outer leaflet, insert into the monolayer that shapes the inner leaflet during the sedimentation of the emulsion droplets. Given the more amphiphilic nature of charged POPA, we anticipate that this insertion is more pronounced for POPA than for POPC. This hypothesis is supported by the data in Figure 1C: asymmetric vesicles with POPC targeted to the inner leaflet exhibit a nonzero αSyn-mEGFP signal when the protein is in the interior of the GUV, suggesting the presence of a small but non-negligible fraction of POPA lipids that must have inserted into the POPC monolayer at the water-oil interface of the emulsion droplets, forming the inner GUV leaflet.

This line of reasoning could potentially provide an interpretation for the stretching elasticity measurements, showing that asymmetric membranes with a POPC:POPA mixture on the outer leaflet display a similar *K*_*A*_ value as symmetric POPC:POPA membranes. With POPA having a higher affinity for the oil-water interface, it has inserted into the monolayer forming the inner leaflet, suppressing the degree of asymmetry and producing a stretching elasticity value similar to that of the symmetric POPC:POPA system (Figure 3C). In the opposite case, where POPC does not compete for insertion into the inner leaflet already containing POPA, it allows for higher asymmetry, resulting in very different stretching elasticity values.

It is not uncommon for asymmetric membranes to exhibit very different mechanical properties compared to their symmetric counterparts. Previous studies have reported increased bending rigidity of asymmetric membranes containing POPC or DOPC in the opposing leaflets compared to symmetric membranes made of the pure counterparts and of the binary mixture [34, 36]. Similar effects were observed for the area compressibility of asymmetric membranes with DMPC or DOPC in the opposing leaflets [37]. The trend of increased bending rigidity of asymmetric vs symmetric membranes was also recently confirmed for asymmetrically charged large unilamellar vesicles containing aminophospholipids [80] (note that in this study, for the asymmetric membranes, only the case of charged lipids present in the inner but not the outer vesicle leaflet was explored). However, to the best of our knowledge, there are no studies reporting stretching elasticity data for asymmetrically charged membranes.

The above results highlight the inherent challenge of accurately predicting the degree of compositional correspondence between the vesicle membrane leaflets and the starting lipid mixtures. This is an important criticism shedding light on the limitations of phase-transfer-based methods in the preparation of asymmetric vesicles. A noteworthy observation is that various previous studies investigating the material properties of asymmetric GUVs tacitly assume that the target leaflet compositions precisely match the starting mixtures. However, due to diverse lipid affinities and partition kinetics at the oil-water interfaces as well as potential lipid and oil mixing, it is plausible that often this assumption may not hold true. In particular, mixtures of lipids with different head groups as examined here can be expected to lead to different final leaflet composition; note that, in a similar way, cholesterol incorporates at much lower fractions than the intended one in vesicles prepared using a phase-transfer method [35].

Finally, we observed an unusual effect of a very small amount of DOPE-based fluorophores on the lateral organization of symmetric POPC:POPA membranes. We noted the presence of gel-like domains that were absent when another dye was employed (Figure 6A-D). Similarly, the presence of the same amount of (label-free) DOPE induced domains in homogeneous membranes (compare Figure 6A and Figure S8G). This outcome could be related to a strong interaction between DOPE and POPA, but we also cannot exclude that it may result from or be enhanced by different dye affinities and partitioning to varying degrees at the oil-water interfaces that form the membrane leaflets. In general, our observations emphasize that particular caution should be exercised when employing fluorescent labels to image and interpret the phase state of vesicles prepared with phase-transfer methods.

The formation of domains was suppressed in the GUVs with asymmetric membranes (Figure 6E and Figure S8H,I). This result aligns with reports emphasizing the dominating effect of the liquid-disordered leaflet on asymmetric membranes [77] as well as the fluidizing effect on ordered phases by asymmetric protein adsorption [81]. These findings, combined with the elasticity results and simulations, underscore the presence of leaflet coupling in asymmetric membranes.

## 5 CONCLUSIONS

The comprehensive series of experiments conducted in this study aimed to elucidate the composition and mechanical properties of symmetric and asymmetric membranes containing POPA. Several key findings have emerged.

The experiments revealed that the outer leaflet composition in vesicles, regardless of the preparation method (electroformation or inverted emulsion), remains consistent within the measurement accuracy. However, characterizing the inner leaflet composition poses challenges due to method limitations, leading to the inference of the presence of POPA without precise quantification.

Stretching elasticity measurements proved to be consistent across different GUV preparation methods for symmetric membranes, suggesting that oil residues, if present, do not substantially alter membrane stretching elasticity. Simulations further supported this, indicating that the stretching elasticity is not a reliable indicator for detecting oil impurities in the membrane. However, simulations did reveal increased membrane thickness and reduced interleaflet coupling in the presence of oil.

Asymmetric membranes displayed similar stretching elasticity values when the outer leaflet was intended to contain a mixture of POPC:POPA and the inner POPC, hinting at a potential insertion of POPA into the inner leaflet during vesicle formation. This interpretation gains support from the conditions of emulsion preparation, where the inner and outer leaflets experience different equilibration times and interface areas, leading to potential asymmetry suppression.

The presence of DOPE-based fluorophores at low fraction (0.5 mol%) was shown to induce gel-like domains in symmetric membranes, cautioning against uncritical use of fluorescent labels in characterizing vesicle phase states. The suppression of these domains in GUVs with asymmetric membranes, underscores the complex interplay of factors influencing the mechanical and compositional properties of asymmetric systems. Overall, this study not only contributes valuable insights into membrane charge asymmetry but also highlights the need for further research and the development of effective characterization techniques for assessing individual leaflet compositions.

## 6 AUTHOR CONTRIBUTIONS

DD, AC and RD proposed and supervised the project. DD and RD designed the experiments. DD, PH, MS and RRMC performed the experiments. DD analyzed the data. DD and RD wrote the manuscript and all authors edited it.

## 8 ACKNOWLEDGEMENTS

This work was possible thanks to the financial support from the National Science Centre (Poland) grant no 2018/30/E/NZ1/00099 (for P.H. and A.C) and Excellence Initiative – Research University for the University of Wrocław (IDUB, Tasks 12 and 3, for D.D. and A.C., respectively). We also thank Agustin Mangiarotti for helping with setting up the micropipette-aspiration setup and Magdalena Zaremba-Czogalla for providing us with the pET28 plasmid containing the mEGFP coding sequence. R.D. thanks M. Miettinen for stimulating discussion.

## REFERENCES

1. Dimova, R., Giant Vesicles and Their Use in Assays for Assessing Membrane Phase State, Curvature, Mechanics, and Electrical Properties.. Annu Rev Biophys, 2019. 48: p. 93–119.

2. Dimova, R. and C. Marques, The Giant Vesicle Book. 2019, Boca Raton: Taylor & Francis Group, LLC.

3. Fenz, S.F. and K. Sengupta, Giant vesicles as cell models. Integrative Biology, 2012. 4(9): p. 982–995.

4. Nair, K.S. and H. Bajaj, Advances in giant unilamellar vesicle preparation techniques and applications. Advances in Colloid and Interface Science, 2023. 318: p. 102935.

5. Zhu, C., et al., Giant Unilamellar Vesicle Microarrays for Cell Function Study. Anal Chem, 2018. 90(24): p. 14363–14367.

6. Gerstle, Z., R. Desai, and S.L. Veatch, Giant Plasma Membrane Vesicles: An Experimental Tool for Probing the Effects of Drugs and Other Conditions on Membrane Domain Stability.. Methods Enzymol, 2018. 603: p. 129–150.

7. Grimmer, M. and K. Bacia, Giant Endoplasmic Reticulum vesicles (GERVs), a novel model membrane tool. Sci Rep, 2020. 10(1): p. 3100.

8. Zhang, W., X. Wang, and X. Han, Multilayer giant unilamellar vesicles as a model of artificial tissue for drug screen. Chemical Physics Letters, 2019. 717: p. 34–37.

9. Kahya, N., Protein-protein and protein-lipid interactions in domain-assembly: lessons from giant unilamellar vesicles. Biochim Biophys Acta, 2010. 1798(7): p. 1392–8.

10. Żelasko, J. and A. Czogalla Selectivity of mTOR-Phosphatidic Acid Interactions Is Driven by Acyl Chain Structure and Cholesterol. Cells, 2022. 11, DOI: 10.3390/cells11010119.

11. Lorent, J.H., et al., Plasma membranes are asymmetric in lipid unsaturation, packing and protein shape. Nat Chem Biol, 2020. 16(6): p. 644–652.

12. van Meer, G., G.W. Voelker Dr Fau - Feigenson, and G.W. Feigenson, Membrane lipids: where they are and how they behave. (1471-0080 (Electronic)).

13. Levental, I. and E. Lyman, Regulation of membrane protein structure and function by their lipid nano-environment. Nature Reviews Molecular Cell Biology, 2022.

14. Doktorova, M., J.L. Symons, and I. Levental, Structural and functional consequences of reversible lipid asymmetry in living membranes. Nat Chem Biol, 2020. 16(12): p. 1321–1330.

15. Zegarlinska, J., et al., Phosphatidic acid - a simple phospholipid with multiple faces. Acta Biochim Pol, 2018. 65(2): p. 163–171.

16. Zhukovsky, M.A., et al., Phosphatidic acid in membrane rearrangements. FEBS Lett, 2019. 593(17): p. 2428–2451.

17. Thakur, R., et al., Regulation of Membrane Turnover by Phosphatidic Acid: Cellular Functions and Disease Implications. Front Cell Dev Biol, 2019. 7: p. 83.

18. Frias, M.A., A. Hatipoglu, and D.A. Foster, Regulation of mTOR by phosphatidic acid. Trends Endocrinol Metab, 2023. 34(3): p. 170–180.

19. Drabik, D. and A. Czogalla, Simple Does Not Mean Trivial: Behavior of Phosphatidic Acid in Lipid Mono- and Bilayers. Int J Mol Sci, 2021. 22(21).

20. Gascard, P., et al., Asymmetric distribution of phosphoinositides and phosphatidic acid in the human erythrocyte membrane. Biochim Biophys Acta, 1991. 1069(1): p. 27–36.

21. Purow, B., Molecular Pathways: Targeting Diacylglycerol Kinase Alpha in Cancer. Clin Cancer Res, 2015. 21(22): p. 5008–12.

22. van Baal, J., et al., Translocation of diacylglycerol kinase theta from cytosol to plasma membrane in response to activation of G protein-coupled receptors and protein kinase C.. J Biol Chem, 2005. 280(11): p. 9870–8.

23. Takaoka, R., et al., Formation of asymmetric vesicles via phospholipase D-mediated transphosphatidylation. Biochim Biophys Acta Biomembr, 2018. 1860(2): p. 245–249.

24. Trauble, H. and E. Grell, Carriers and specificity in membranes. IV. Model vesicles and membranes. The formation of asymmetrical spherical lecithin vesicles.. Neurosci Res Program Bull, 1971. 9(3): p. 373–80.

25. Szoka, F., Jr. and D. Papahadjopoulos, Comparative properties and methods of preparation of lipid vesicles (liposomes). Annu Rev Biophys Bioeng, 1980. 9: p. 467–508.

26. Peyret, A., H. Zhao, and S. Lecommandoux, Preparation and Properties of Asymmetric Synthetic Membranes Based on Lipid and Polymer Self-Assembly. Langmuir, 2018. 34(11): p. 3376–3385.

27. Huang, Y., S.H. Kim, and L.R. Arriaga, Emulsion templated vesicles with symmetric or asymmetric membranes. Adv Colloid Interface Sci, 2017. 247: p. 413–425.

28. Pautot, S., B.J. Frisken, and D.A. Weitz, Engineering asymmetric vesicles. Proc Natl Acad Sci U S A, 2003. 100(19): p. 10718–21.

29. Noireaux, V. and A. Libchaber, A vesicle bioreactor as a step toward an artificial cell assembly. Proc Natl Acad Sci U S A, 2004. 101(51): p. 17669–74.

30. Enoki, T.A., et al., Dataset of asymmetric giant unilamellar vesicles prepared via hemifusion: Observation of anti-alignment of domains and modulated phases in asymmetric bilayers.. Data Brief, 2021. 35: p. 106927.

31. Scott, H.A.-O., et al., Model Membrane Systems Used to Study Plasma Membrane Lipid Asymmetry. Symmetry, 2021. 13(8): p. 1356.

32. Doktorova, M., et al., Preparation of asymmetric phospholipid vesicles for use as cell membrane models. Nat Protoc, 2018. 13(9): p. 2086–2101.

33. Richmond, D.L., et al., Forming giant vesicles with controlled membrane composition, asymmetry, and contents. Proc Natl Acad Sci U S A, 2011. 108(23): p. 9431–6.

34. Karamdad, K., et al., Studying the effects of asymmetry on the bending rigidity of lipid membranes formed by microfluidics. Chem Commun (Camb), 2016. 52(30): p. 5277–80.

35. Blosser, M.C., B.G. Horst, and S.L. Keller, cDICE method produces giant lipid vesicles under physiological conditions of charged lipids and ionic solutions. Soft Matter, 2016. 12(35): p. 7364–7371.

36. Elani, Y., et al., Measurements of the effect of membrane asymmetry on the mechanical properties of lipid bilayers. Chem Commun (Camb), 2015. 51(32): p. 6976–9.

37. Lu, L., et al., Membrane mechanical properties of synthetic asymmetric phospholipid vesicles. Soft Matter, 2016. 12(36): p. 7521–7528.

38. Leomil, F.S.C., et al., Bilayer Charge Asymmetry and Oil Residues Destabilize Membranes upon Poration. Langmuir, 2024.

39. Mizuno, S., et al., Dioleoyl-phosphatidic acid selectively binds to alpha-synuclein and strongly induces its aggregation. FEBS Lett, 2017. 591(5): p. 784–791.

40. Yamada, H., et al., Characterization of alpha-synuclein N-terminal domain as a novel cellular phosphatidic acid sensor. FEBS J, 2020. 287(11): p. 2212–2234.

41. Moga, A., et al., Optimization of the Inverted Emulsion Method for High-Yield Production of Biomimetic Giant Unilamellar Vesicles. Chembiochem, 2019. 20(20): p. 2674–2682.

42. Stephan, M.S., et al., Biomimetic asymmetric bacterial membranes incorporating lipopolysaccharides. Biophys J, 2022.

43. Pautot, S., B.J. Frisken, and D.A. Weitz, Production of unilamellar vesicles using an inverted emulsion. Langmuir, 2003. 19(7): p. 2870–2879.

44. Rouser G Fau - Fkeischer, S., A. Fkeischer S Fau - Yamamoto, and A. Yamamoto, Two dimensional then layer chromatographic separation of polar lipids and determination of phospholipids by phosphorus analysis of spots.. Lipids, 1970(5): p. 494–496.

45. Hope, M.J., et al., Generation of multilamellar and unilamellar phospholipid vesicles. Chemistry and Physics of Lipids, 1986. 40(2): p. 89–107.

46. van den Ent, F. and J. Lowe, RF cloning: a restriction-free method for inserting target genes into plasmids. J Biochem Biophys Methods, 2006. 67(1): p. 67–74.

47. Laemmli, U.K., Cleavage of structural proteins during the assembly of the head of bacteriophage T4. Nature, 1970. 227(5259): p. 680–5.

48. Steinkuhler, J., et al., Charged giant unilamellar vesicles prepared by electroformation exhibit nanotubes and transbilayer lipid asymmetry. Sci Rep, 2018. 8(1): p. 11838.

49. Carvalho, K., et al., Giant unilamellar vesicles containing phosphatidylinositol(4,5)bisphosphate: characterization and functionality. Biophys J, 2008. 95(9): p. 4348–60.

50. Philips, J.C., et al., Scalable molecular dynamics with NAMD. J Comput Chem, 2005. 26(16): p. 1781–1802.

51. Klauda, J.B., et al., Update of the CHARMM all-atom additive force field for lipids: validation on six lipid types. J Phys Chem B, 2010. 114(23): p. 7830–43.

52. Jo, S., et al., CHARMM-GUI: a web-based graphical user interface for CHARMM. J Comput Chem, 2008. 29(11): p. 1859–65.

53. Vanommeslaeghe, K., et al., CHARMM general force field: A force field for drug-like molecules compatible with the CHARMM all-atom additive biological force fields.. J Comput Chem, 2010. 31(4): p. 671–90.

54. Lee, J., et al., CHARMM-GUI Input Generator for NAMD, GROMACS, AMBER, OpenMM, and CHARMM/OpenMM Simulations Using the CHARMM36 Additive Force Field.. J Chem Theory Comput, 2016. 12(1): p. 405–13.

55. Giorgino, T., Computing 1-D atomic densities in macromolecular simulations: The density profile tool for VMD. Computer Physics Communications, 2014. 185(1): p. 317–322.

56. Doktorova, M., et al., A New Computational Method for Membrane Compressibility: Bilayer Mechanical Thickness Revisited. Biophys J, 2019. 116(3): p. 487–502.

57. Guixa-Gonzalez, R., et al., MEMBPLUGIN: studying membrane complexity in VMD. Bioinformatics, 2014. 30(10): p. 1478–80.

58. Tarun, O.B., M.Y. Eremchev, and S. Roke, Interaction of Oil and Lipids in Freestanding Lipid Bilayer Membranes Studied with Label-Free High-Throughput Wide-Field Second-Harmonic Microscopy.. Langmuir, 2018. 34(38): p. 11305–11310.

59. Stano, P., Commentary: Rapid and facile preparation of giant vesicles by the droplet transfer method for artificial cell construction.. Front Bioeng Biotechnol, 2022. 10: p. 1037809.

60. Vitkova, V., J. Genova, and I. Bivas, Permeability and the hidden area of lipid bilayers. European Biophysics Journal with Biophysics Letters, 2004. 33(8): p. 706–714.

61. Drabik, D., et al., Determination of the Mechanical Properties of Model Lipid Bilayers Using Atomic Force Microscopy Indentation. Langmuir, 2020. 36(44): p. 13251–13262.

62. Drabik, D., et al., Mechanical Properties Determination of DMPC, DPPC, DSPC, and HSPC Solid-Ordered Bilayers. Langmuir, 2020. 36(14): p. 3826–3835.

63. Faizi, H.A., et al., Bending Rigidity, Capacitance, and Shear Viscosity of Giant Vesicle Membranes Prepared by Spontaneous Swelling, Electroformation, Gel-Assisted, and Phase Transfer Methods: A Comparative Study.. Langmuir, 2022. 38(34): p. 10548–10557.

64. Park, S., W. Im, and R.W. Pastor, Developing initial conditions for simulations of asymmetric membranes: a practical recommendation. Biophys J, 2021. 120(22): p. 5041–5059.

65. Chaisson, E.H., F.A. Heberle, and M. Doktorova Building Asymmetric Lipid Bilayers for Molecular Dynamics Simulations: What Methods Exist and How to Choose One. Membranes, 2023. 13, DOI: 10.3390/membranes13070629.

66. Miettinen, M.S. and R. Lipowsky, Bilayer Membranes with Frequent Flip-Flops Have Tensionless Leaflets. Nano Letters, 2019. 19(8): p. 5011–5016.

67. Doktorova, M. and H. Weinstein, Accurate In Silico Modeling of Asymmetric Bilayers Based on Biophysical Principles. Biophysical Journal, 2018. 115(9): p. 1638–1643.

68. Rucker, G., X. Yu, and L.Q. Zhang, Molecular dynamics investigation on n-alkane-air/water interfaces. Fuel, 2020. 267.

69. Nagle, J.F., Area Compressibility Moduli of the Monolayer Leaflets of Asymmetric Bilayers from Simulations. Biophys J, 2019. 117(6): p. 1051–1056.

70. Rawicz, W., et al., Effect of chain length and unsaturation on elasticity of lipid bilayers. Biophys J, 2000. 79(1): p. 328–39.

71. Evans, E.A., Bending resistance and chemically induced moments in membrane bilayers. Biophys J, 1974. 14(12): p. 923–31.

72. Goetz, R., G. Gompper, and R. Lipowsky, Mobility and Elasticity of Self-Assembled Membranes. Physical Review Letters, 1999. 82(1): p. 221–224.

73. Demel, R.A., et al., Monolayer characterstics and thermal behaviour of phosphatidic acids. Chemistry and Physics of Lipids, 1992. 60(3): p. 209–223.

74. Kuppe, K., et al., Calcium-Induced Membrane Microdomains Trigger Plant Phospholipase D Activity. ChemBioChem, 2008. 9(17): p. 2853–2859.

75. Kooijman, E.E., et al., What Makes the Bioactive Lipids Phosphatidic Acid and Lysophosphatidic Acid So Special. Biochemistry, 2005. 44(51): p. 17007–17015.

76. Kooijman, E.E., et al., An Electrostatic/Hydrogen Bond Switch as the Basis for the Specific Interaction of Phosphatidic Acid with Proteins*.. Journal of Biological Chemistry, 2007. 282(15): p. 11356–11364.

77. Arribas Perez, M. and P.A. Beales, Dynamics of asymmetric membranes and interleaflet coupling as intermediates in membrane fusion. Biophysical Journal, 2023. 122(11): p. 1985–1995.

78. Enoki, T.A. and F.A. Heberle, Experimentally determined leaflet–leaflet phase diagram of an asymmetric lipid bilayer. Proceedings of the National Academy of Sciences, 2023. 120(46): p. e2308723120.

79. Lin, Q. and E. London, Ordered Raft Domains Induced by Outer Leaflet Sphingomyelin in Cholesterol-Rich Asymmetric Vesicles. Biophysical Journal, 2015. 108(9): p. 2212–2222.

80. Frewein, M.P.K., et al., Distributing aminophospholipids asymmetrically across leaflets causes anomalous membrane stiffening. Biophysical Journal, 2023. 122(12): p. 2445–2455.

81. Pataraia, S., et al., Effect of cytochrome c on the phase behavior of charged multicomponent lipid membranes. Biochimica Et Biophysica Acta - Biomembranes, 2014. 1838(8): p. 2036–2045.

