## Supporting Information for "Effect of leaflet asymmetry on the stretching elasticity of lipid bilayers with phosphatidic acid"

### 1. Sequence of primers in cloning procedure

**Table S1.** Sequence of primers used in the cloning procedures.

| Primer name | 5' → 3' sequence | Purpose of use |
|---|---|---|
| **aS-N BamH Fwd** | TACAGGATCCATGGATGTGTTCATGAAAGGAC | Amplification of N-terminal α-syn fragment and addition of restriction digest sites for BamHI and EcoRI enzymes |
| **aS-N EcoRI Rev** | ATACGAATTCGTTTGGTCTTCTCAGCCACTG | |
| **aS Q61-A140 Fwd** | GACAACAGTGGCTGAGAAGACCAAAGAGCAAGTGACAAATGTTGGAG | Restriction free cloning of Q61-A140 α-syn fragment into plasmid containing M1-K60 fragment to obtain construct coding full length α-syn |
| **aS Q61-A140 Rev** | CCCTGAAACAGCACTTCCAGAATTCAGGCTTCAGGCTCATAGTCTTG | |
| **Δ STOP Fwd** | ATTCTGGAAGTGCTGTTTC | Mutagenic primers used to remove the stop codon located between the sequence encoding α-synuclein and the mEGFP protein and obtain construct encoding α-synuclein with fused mEGFP protein |
| **Δ STOP Rev** | GGCTTCAGGCTCATAGTC | |
| **T7 promoter** | TAATACGACTCACTATAGGG | Sequencing of coding sequence between T7 promoter and T7 terminator in obtained DNA constructs |
| **T7 terminator** | TGCTAGTTATTGCTCAGCGG | |

### 2. SDS-PAGE analysis of purified proteins

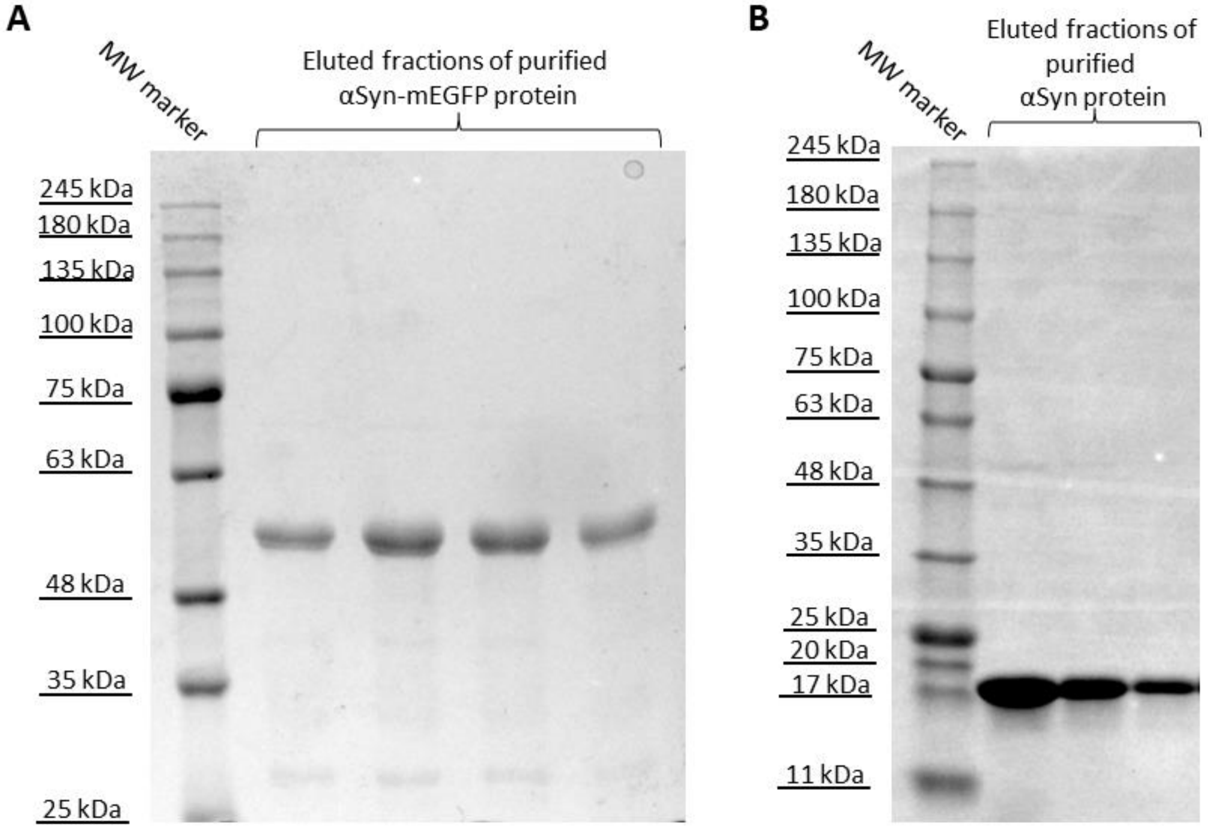


**Figure S1**. SDS-PAGE analysis of purified recombined αSyn-mEGFP (A) and αSyn (B) proteins.

### 3. Optimization of αSyn-mEGFP concentration for fluorescence quantification

In order to identify the minimal suitable concentration of αSyn-mEGFP for microscopy observations of GUV containing PA, three different protein concentrations were tested on electroformed vesicles – 100 nM, 250 nM and 500 nM. Both 500 nM and 250 nM resulted in sufficient signal with the used microscope settings (Figure 2SA). The same settings were later used to assess the PA content in inverted-emulsion vesicles. Note that to be able to apply the findings for suitable protein concentration optimized on electroformed vesicles, we ensured that the lipid concentrations in the GUV samples, and thus the lipid-protein ratio, were similar for both GUV preparation methods (see SI section 8). It is interesting to emphasize that the amount of lipids needed to prepare electroformed GUVs at lipid concentration similar to that for GUVs prepared with the emulsion transfer is roughly five-fold lower.

Additionally, the effect of NaCl was investigated. When no NaCl was added, a small but significant decrease of the intensity was observed for vesicles with 500 nM protein, and no significant change for vesicles with 250 nM protein (Figure 2SA). Because the signal was sufficiently strong, the evaluation of the inverted-emulsion vesicles was conducted in the absence of salt. The change of intensity was also investigated as a function of PA fraction in the membrane and the trend was found nonlinear, see Figures S2B and S3. For membranes containing 5 and 10 mol% PA, the signal is indistinguishable and close to that of PA-free membranes pointing to limitations of the method.

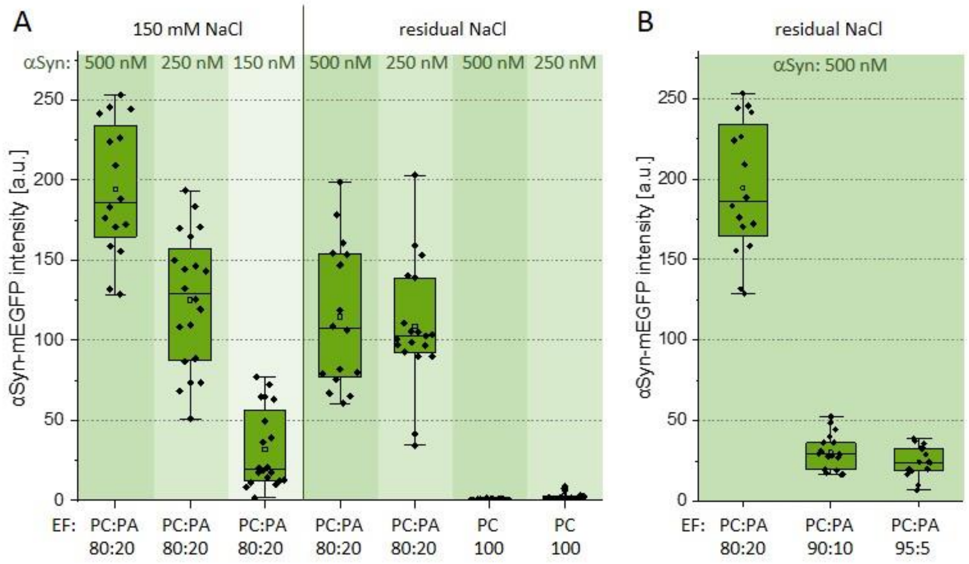


**Figure S2.** Fluorescence intensity of αSyn-mEGFP measured on electroformed (EF, symmetric) vesicles at different conditions. (A) Intensity signal on the vesicle membrane as a function of bulk protein concentration in the presence of 150 mM NaCl or residual amounts of NaCl (introduced with the protein buffer solutions roughly corresponding to 1 mM NaCl final concentration in the GUV suspension). For comparison, data with POPA-free membranes (pure POPC) are also shown. (B) Intensity as a function of PA fraction in POPC:POPA vesicles in the presence of 500 nM αSyn-mEGFP and residual NaCl (corresponding to roughly 1 mM NaCl).

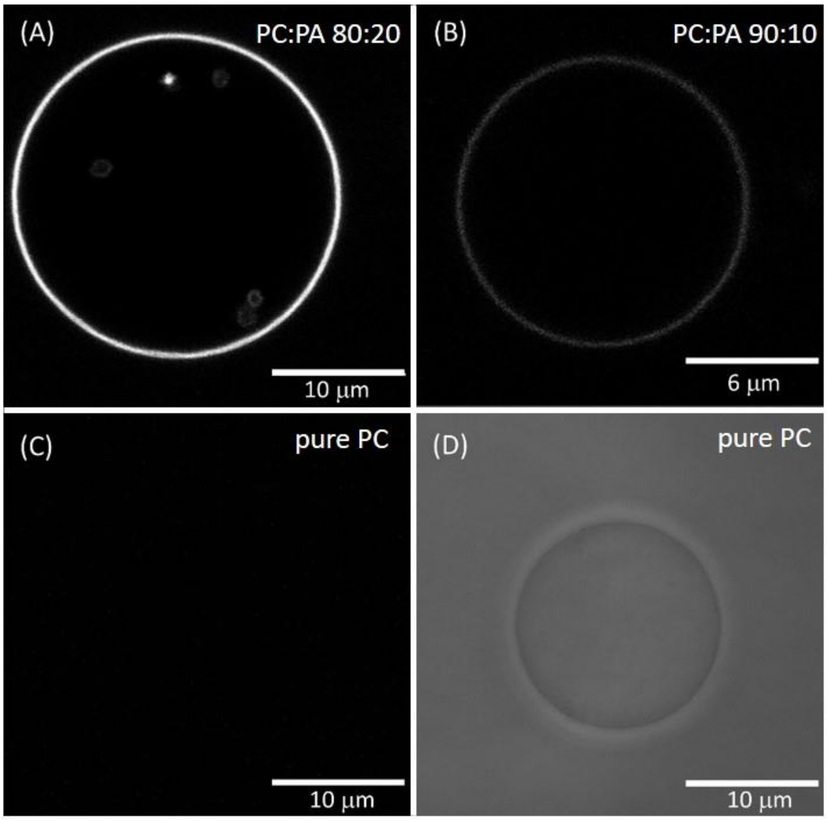


**Figure S3.** Snapshots of electroformed (symmetric) POPC:POPA vesicles containing different PA fraction and in the presence of 0.5 μM αSyn-mEGFP and residual NaCl. The fluorescence signal in (A-C) is from αSyn-mEGFP. (A) POPC:POPA 80:20; (B) POPC:POPA 90:10; (C) POPC; (D) The same POPC vesicle as in panel C observed under phase contrast.

### 4. Zeta-potential measurements on GUVs

Zeta potential on GUVs was measured in three different solutions: without salt, with salt (as presented in the main text) and in a buffer: First, we used the procedure available in the literature [1, 2]. Specifically, U-cells (Malvern DTS1070) of volume 1 mL were used with voltage for electrophoretic movement set to 150 V. These results are presented in Figure S4A. The zeta-potential of symmetric and asymmetric vesicles with external leaflet made of POPC and those with POPC:POPA 80:20 as external leaflet showed a relatively small difference (although statistically significant). Overall, there was also an issue with the measurements of the inverted-emulsion vesicles, namely that insufficiently high number of objects were present in the sample, which resulted in poor quality report and rejection. We thus implemented two changes. To reduce the dilution of the GUV suspensions required for the large-volume U-cells, we employed dip-cells (Malvern ZEN1002), which require only 0.6 mL of sample. Secondly, we added 5 mM NaCl to the external solution to increase the salinity as advised by the manufacturer (for example, to reduce issues associated with electrode polarization). Having used different cell with a different path between the electrodes, the voltage for electrophoretic movement had to be adjusted and was set to 10 V. The results are presented in Figure 2 in the main text showing substantial difference between the zeta potential of POPC and POPC:POPA 80:20 in the outer leaflets. Finally, because PA is a lipid with protonation sensitive to pH, we performed measurements in 10 mM HEPES buffer of pH 7.4 to ensure that results are similar in buffered environment. The obtained zeta potentials for conditions in the absence of salt and in the presence of buffer are presented in Figure S4B. The difference between vesicles with PC vs PC:PA lipids in the outer leaflet was preserved, but altered in magnitude because of the low conductivity of the solution (compare to Figure 2 in the main text). Finally, no substantial changes in the zeta potential were detected to change over time (Figure S4C) implying no significant interleaflet exchange.

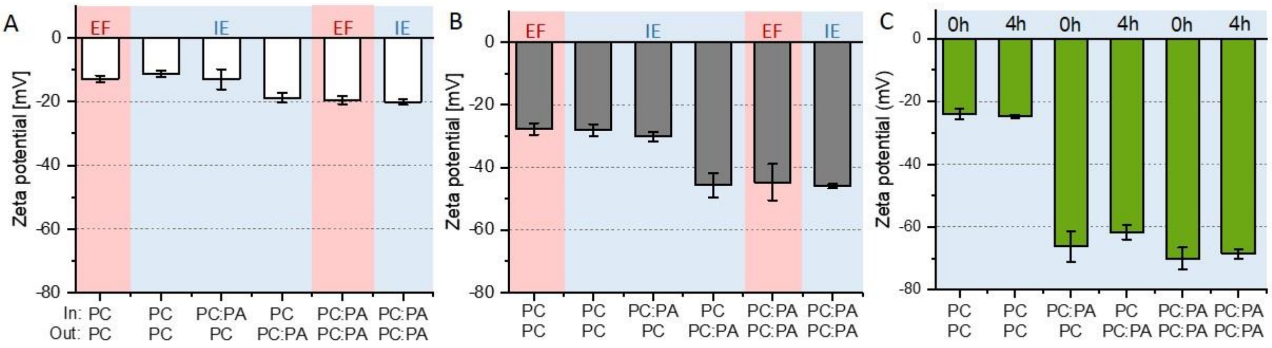


**Figure S4.** Zeta-potential values of GUVs measured in solutions of different salinity and over time. Internal and external leaflets (labelled as "In" and "Out") with composition POPC:POPA 80:20 (molar ratio) are indicated as PC:PA, and pure POPC as PC. Samples prepared with the inverted emulsion method (IE) are shaded in light blue, and those with electroformation (EF) – in pink. The vesicle were in (A) sucrose/glucose solutions, (B) sucrose/glucose solutions buffered with 10 mM HEPES, pH 7.4, and (C) in sucrose/glucose solution containing 5 mM NaCl (as in the main text) right after preparation and after 4 hours; the zeta potential data measured immediately after preparation (0h) is the same as that in Figure 2A in the main text. The measurements were performed at 25 °C.

### 5. Micropipette aspiration data

The vesicle-micropipette measurements were first optimized and probed for hysteresis potentially resulting from membrane adhesion to the glass capillaries. Individual results of micropipette aspiration experiments are presented in Figure S5 for symmetric and asymmetric vesicles prepared via electroformation and inverted emulsion transfer.

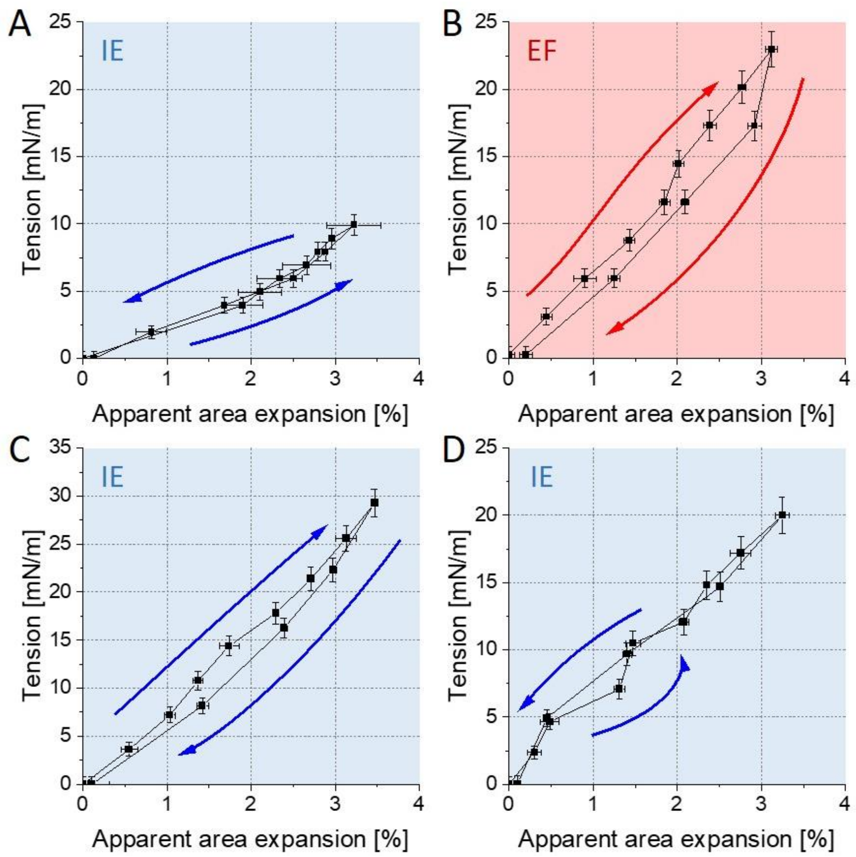


**Figure S5**. Example experimental data testing for hysteresis for both electroformed (pink background) and inverted-emulsion (light blue) vesicles composed of (A) symmetric, POPC, (B) symmetric, POPC:POPA 80:20, (C) asymmetric, Out: POPC:POPA, In: POPC and (D) asymmetric, Out POPC, In: POPC:POPA.

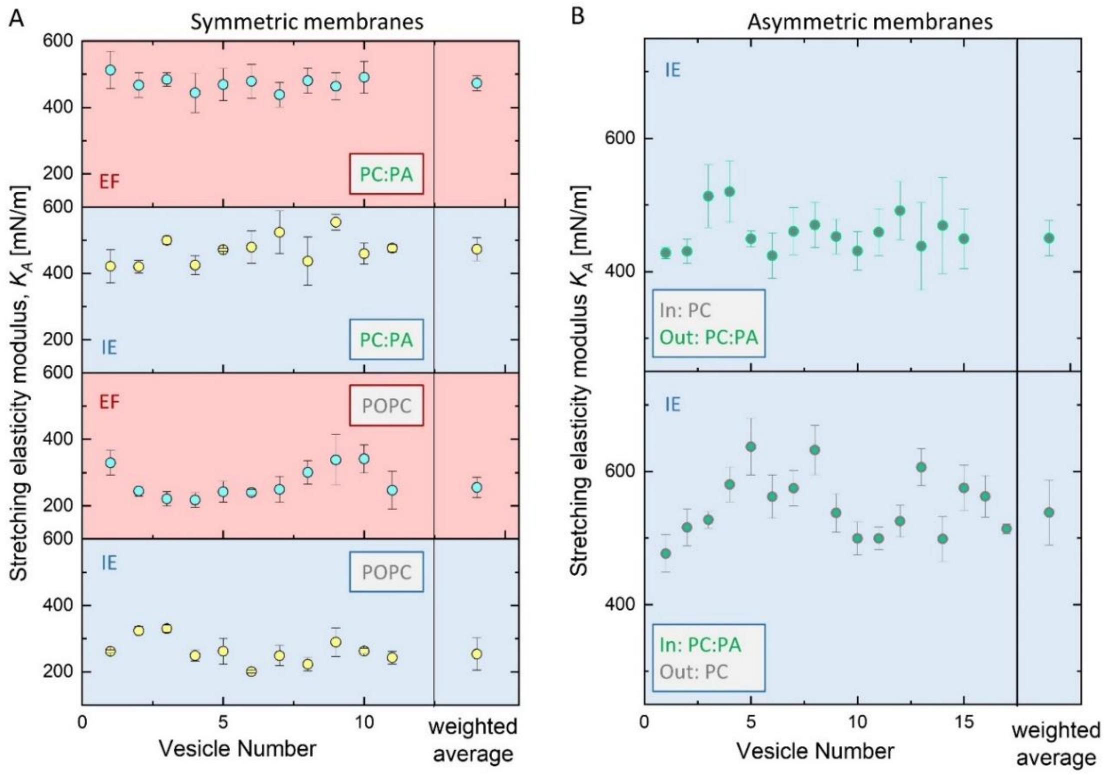


**Figure S6.** Area compressibility values measured on individual GUVs with (A) symmetric and (B) asymmetric membranes prepared using electroformation (EF, pink background) or inverted-emulsion method (IE, light blue). The values plotted on the right represents weighted area compressibility, where the weight was assumed as inverse of given measurement uncertainty.

### 6. Differential scanning calorimetry data

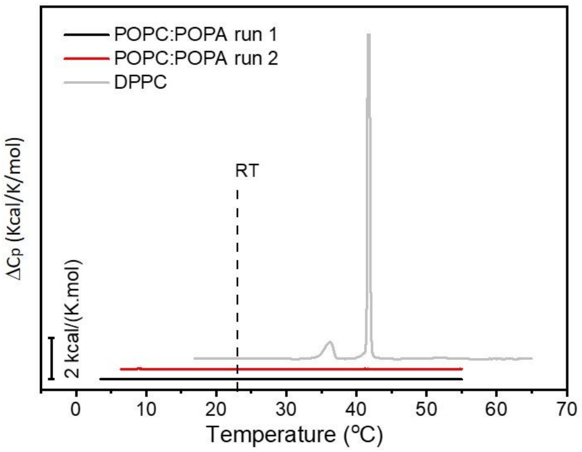


**Figure S7.** Excess heat capacity profiles of LUVs composed of POPC:POPA 80:20 (black and red, 5 mM total lipid concentration) show no detectable phase transition; for comparison a measurement on MLVs composed of dipalmitoylphosphatidylcholine (DPPC, 10 mM lipid concentration) exhibiting a gel-fluid phase transition at 41°C is shown in gray. Room temperature (RT) is indicated by a vertical black dashed line. For the POPC:POPA mixture, the temperature ranges for run 1 and run 2 were 5-55 °C and 2-55 °C respectively, at a scan rate of 20°C/h (heating). For the DPPC composition, the temperature range was 15-65 °C at a scan rate of 60°C/h (heating). All scans were normalized to the respective lipid concentration. The curves were shifted in the y-axis for better visualization.

### 7. Phase separation induced by DOPE-based dyes

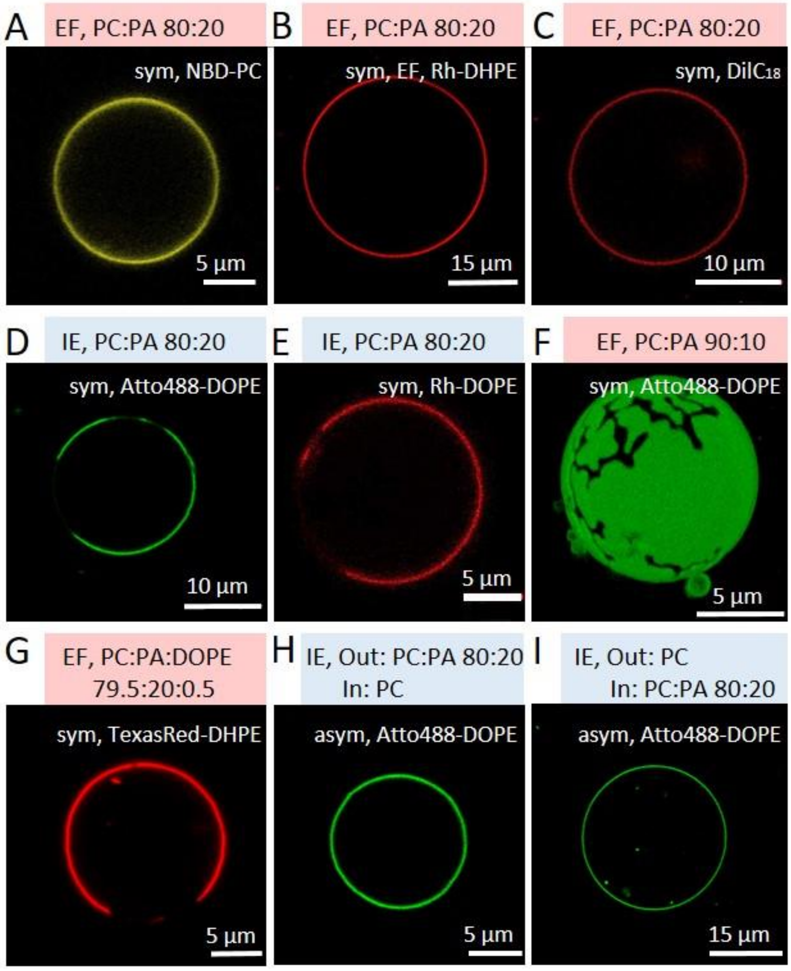


**Figure S8.** Effect of membrane composition and fluorescent label (0.5 mol%) on domain appearance. Examples of symmetric POPC:POPA 80:20 electroformed (EF, pink highlight) vesicles labelled with (A) NBD-PC, (B) Rh-DHPE and (C) $DilC_{18}$, not showing phase separation; for clarity we show confocal cross sections even though the whole vesicle surface was examined for domains. Examples of POPC:POPA 80:20 inverted-emulsion (IE, light blue) vesicles labelled with (D) Atto488-DOPE and (E) Rh-DOPE showing phase separation; the dyes were added to both oil phases forming the individual leaflets. (F) EF GUV made of POPC:POPA 90:10 labeled with Atto488-DOPE showing smaller domain area fraction compared to 80:20. (G) EF GUV made of POPC:POPA:DOPE 79.5:20:0.5 labelled with TexasRed-DHPE. Examples of asymmetric IE vesicles without domains: (H) POPA-containing outer leaflet labelled with Atto488-DOPE and (I) POPA-containing inner leaflet labelled with Atto488-DOPE.

### 8. Assessment of lipid amount in the GUV samples and the efficiency of the electroformation and inverted emulsion preparation methods in terms of used lipids

The assessment of lipid amount in the GUV samples was performed with phosphorus analysis [3]. Ascorbic acid was bought from Roth, ammonium heptamolybdate was bought from POCH, Phosphorus Standard for AAS was purchased from Fluka Analytical. A calibration curve (Figure S9A) was built as follows. Appropriate amounts of $KH_2PO_4$ that correspond to 0, 0.5, 1, 0.5, 2, 2.5 and 5 µg of $KH_2PO_4$ were dissolved in 50 µl of water in glass vials. This was followed by addition of 0.5 ml of concentrated perchloric acid (70%) to all vials and heating to 200 °C for 2 hours under cover to mineralize phosphate. Then, 1ml aqueous solution of 2.5 wt% ammonium molybdate and 10 wt% ascorbic acid was added to each vial, followed by vortexing and keeping the samples for 1 hour under 37 °C. The samples were then transferred to 96-well plate in such a way that each sample was pipetted in at least 3 wells. After cooling, the samples absorbance was measured at 800 nm wavelength using Asys UVM340 Plate Reader (Biochrom). The GUV samples (1.5 ml) were evaporated overnight under 95 °C and re-suspended in 50 µl water to ensure similar conditions as for the samples used for preparing a calibration curve. They were subsequently treated as described above. The lipid mass was estimated from the calibration curve and the measured absorbance of the GUV samples, see Figure S9B. The efficiency of the preparation methods was calculated as the ratio of total amount of lipid in the obtained GUVs sample to the total amount of lipid applied to the electrode (in the electroformation protocol) or added to the two oil phases (in the inverted emulsion method).

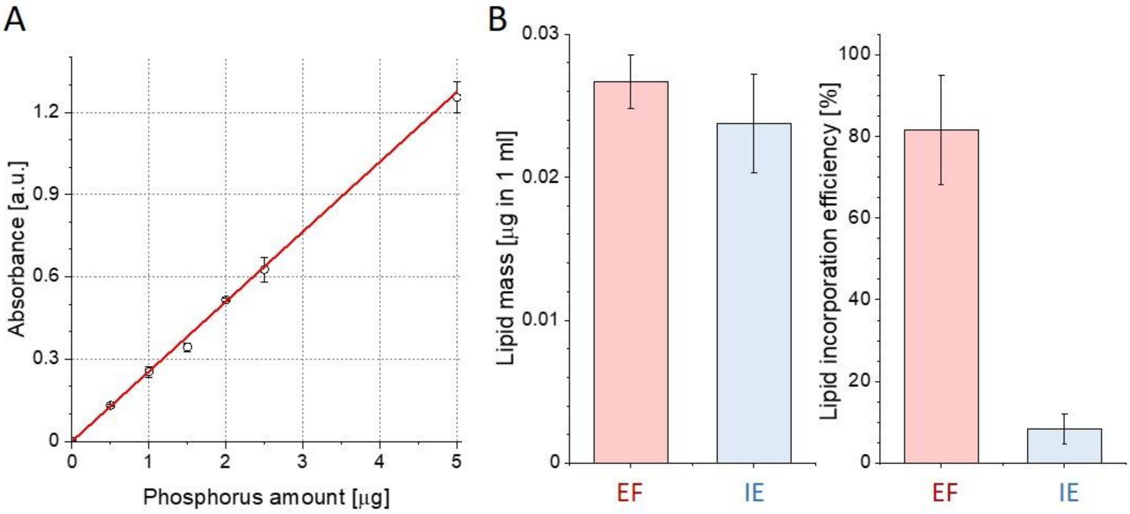


**Figure S9.** (A) Calibration curve used to determine lipid mass in GUV solutions (see text for details). (B) Lipid mass in 1 ml solutions of GUV suspensions obtained via electroformation (EF) or inverted emulsion (IE) method (left) and efficiency of lipid incorporation calculated as lipid amount in the GUV sample vs the initial amount of lipid used for the preparation for symmetric POPC vesicles (right).

### References

1. Steinkuhler, J., et al., *Charged giant unilamellar vesicles prepared by electroformation exhibit nanotubes and transbilayer lipid asymmetry.* Sci Rep, 2018. **8**(1): p. 11838.
2. Carvalho, K., et al., *Giant unilamellar vesicles containing phosphatidylinositol(4,5)bisphosphate: characterization and functionality.* Biophys J, 2008. **95**(9): p. 4348-60.
3. Rouser G Fau - Fkeischer, S., A. Fkeischer S Fau - Yamamoto, and A. Yamamoto, *Two dimensional then layer chromatographic separation of polar lipids and determination of phospholipids by phosphorus analysis of spots.* Lipids, 1970(5): p. 494-496.
